# HIF-1α-mediated mitochondrial-glycolytic reprogramming controls the transition of precursor to terminally exhausted T cells

**DOI:** 10.1101/2023.08.31.555662

**Authors:** Hao Wu, Xiufeng Zhao, Sophia M. Hochrein, Miriam Eckstein, Gabriela F. Gubert, Konrad Knöpper, Ana Maria Mansilla, Arman Öner, Remi Doucet-Ladevèze, Werner Schmitz, Bart Ghesquière, Sebastian Theurich, Jan Dudek, Georg Gasteiger, Alma Zernecke-Madsen, Sebastian Kobold, Wolfgang Kastenmüller, Martin Vaeth

## Abstract

Functional exhaustion of T cells in cancer and persistent infections is characterized by the upregulation of inhibitory receptors, the progressive decline in cytokine secretion and impaired cytolytic activity. Terminally exhausted T cells are steadily replenished by a precursor population (Tpex) with phenotypic features of memory T cells and a stem-like capacity to self-renew. However, the metabolic principles of Tpex maintenance and the regulatory circuits that control the exhaustion of their progeny remain incompletely understood. Using a combination of gene-deficient mice, single-cell transcriptomics and metabolomic analyses, we here show that mitochondrial insufficiency is a cell-intrinsic trigger that initiates the T cell exhaustion program. At the molecular level, we found that diminished mitochondrial respiration and metabolic remodeling cause oxidative stress, which inhibits the proteasomal degradation of *hypoxia inducible factor 1 alpha* (HIF-1α) in Tpex cells. HIF-1α mediates the transcriptional-glycolytic reprogramming of Tpex cells as an initial step towards terminal differentiation and functional exhaustion. Finally, we show that enhancing respiration by limiting the glycolytic activity of CAR T cells is a feasible metabolic intervention strategy to preserve the stemness of Tpex cells during chronic viral infection and cancer immunotherapy.

---

Following an acute infection, cognate CD8^+^ T cells proliferate and differentiate into cytotoxic T lymphocytes (CTLs) to eliminate infected or transformed target cells. The differentiation of CTLs is paralleled by transcriptional, epigenetic and metabolic rewiring to support their clonal expansion and effector functions, such as cytokine secretion and cytotoxicity ^1, 2, 3^. After the clearance of the pathogen and the resolution of inflammation, most effector T cells die and only a few lymphocytes remain as long-lived memory T cells ^3^.

However, when (self-) antigens cannot be eliminated, such as in persistent infection and cancer, continuous antigen receptor stimulation drives an alternative differentiation trajectory, known as T cell exhaustion ^1, 2, 4^. Exhausted T (Tex) cells are characterized phenotypically by the upregulation of co-inhibitory receptors, such as PD-1, Tim-3 and Lag-3, and functionally by the decline in cytokine secretion and cytolytic activity ^1, 4^. Although Tex cells also lose proliferative capacity and undergo apoptosis during terminal differentiation, they are steadily replenished by a distinct multipotent population, termed precursors (or progenitors) of exhausted T (Tpex) cells ^4, 5, 6, 7, 8^. Tpex cells can be identified by the expression of memory-related molecules, such as the transcription factor TCF-1 and the cell surface molecules Ly108 and CD62L, but the absence of terminal differentiation markers, such as Tim-3, CD101 or 2B4 ^4, 5, 8, 9, 10, 11^. The ‘stem-like’ capacity of Tpex cells to self-renew and their superior repopulation potential following adoptive transfer into chronically infected mice spurred an increasing scientific interest to define the molecular mechanisms that control the transition from multipotent Tpex into dysfunctional Tex cells ^4, 7^. This question is clinically relevant, since Tpex cells are the main mediators of the antitumor immune responses following therapeutic checkpoint blockade ^4, 6, 7, 9, 12^. Although PD-1 expression is found on both Tpex and terminally exhausted T cells, only Tpex cells respond efficiently to anti-PD-1 treatment with a proliferative burst and the differentiation into cytotoxic T cells, thus reinvigorating ‘exhausted’ antitumor immune responses in patients following checkpoint immunotherapy ^4, 6, 7, 9, 12, 13, 14^. However, the molecular mechanisms that maintain the stemness of Tpex cells and the signals that instruct their functional exhaustion remain incompletely understood and merit further investigation to improve the clinical outcome of cancer immunotherapy.

Naïve T cells are characterized by a low-rate catabolic metabolism and generate ATP primarily through mitochondrial respiration and fatty acid oxidation (FAO) ^15, 16^. Antigen receptor ligation triggers a Ca^2+^ influx cascade in T cells, which activates the serine/threonine phosphatase calcineurin promoting the nuclear translocation of *nuclear factor of activated T cells* (NFAT) ^17,18^. In cooperation with other transcription factors, NFAT controls the expression of nutrient transporters and metabolic enzymes to support the cellular growth and clonal expansion of activated T cells ^19, 20, 21^. To meet their increased bioenergetic demand, activated T cells rewire their metabolic machinery and utilize aerobic glycolysis and oxidative phosphorylation (OXPHOS) for ATP production and the biosynthesis of building blocks ^16, 19, 20^. Whereas most effector T cells perish following antigen clearance, a small fraction persists as memory T cells, which rely on mitochondrial respiration.

Although metabolic reprogramming controls multiple aspects of T cell activation and effector differentiation ^15, 16, 22^, the regulation of cellular metabolism and its contribution to terminal T cell exhaustion remains incompletely understood ^2^. Recent studies have demonstrated that mitochondrial decline correlates with T cell exhaustion *in vitro* ^23, 24^, during chronic viral infection ^25, 26^ and in cancer ^27, 28, 29^, but whether these changes are cause or consequence of T cell exhaustion remained unclear. We here provide genetic evidence that impaired mitochondrial respiration is not merely a consequence of T cell dysfunction but, instead, sufficient to elicit the transcriptional, phenotypic and functional hallmarks of T cell exhaustion. Mechanistically, we found that oxidative stress due to mitochondrial deterioration and metabolic rewiring antagonizes the proteasomal degradation of *hypoxia-inducible factor 1* α (HIF-1α), which mediates the glycolytic reprogramming of Tpex cells as an initial step towards terminal differentiation. Our findings also demonstrate that mitochondrial respiration is a prerequisite for the stemness of exhausted T cells and that limiting their glycolytic capcaity is a promising metabolic strategy to maintain the functionality of virus-specific and *chimeric antigen receptor* (CAR) T cells during chronic viral infection and cancer.

## RESULTS

### Terminal T cell exhaustion is characterized by metabolic reprogramming

To analyze the transcriptional-metabolic programs alongside the developmental trajectories of exhausted T cells, we performed single-cell (sc) RNA sequencing of CD44^+^ PD-1^hi^ CD8^+^ T cells isolated from chronically infected wild-type (WT) mice after infection with the ‘clone 13’ strain of lymphocytic choriomeningitis virus (LCMV^CL13^) (Extended Data Fig. 1a). To avoid a bias due to active cell division, we focused on non-proliferating T cells and identified six major populations of exhausted T cells using unsupervised clustering visualized by uniform manifold approximation and projection (UMAP) (Fig. 1a and Extended Data Fig. 1b). Tpex cells were characterized by a unique combination of activation (*Cd44, Cd69, Icos*), exhaustion (*Pdcd1*, *Tox*) and memory-related molecules (*Sell, Ccr7, Tcf7, Slamf6, Id3*, *Cxcr5*) (Fig. 1b-d and Extended Data Fig. 1b,c). Diffusion pseudotime analysis predicted two differentiation trajectories originating from ‘stem-like’ CD62L^+^ Tpex1 cells ^10^ (Fig. 1b). One trajectory merged into a ‘CTL-like’ exhausted T cell population that is characterized by *Cx3cr1*, *Klrg1* and *S100a4* expression (Extended Data Fig. 1c) ^30^. The second lineage pointed towards terminally exhausted T cells (Fig. 1b). Indeed, the Tex1 and Tex2 subsets showed a progressive upregulation of terminal exhaustion markers, such as *Havcr2* (encoding for Tim-3), *Cxcr6*, *Id2*, *Lag3*, *Gzma*, *Cd244a* (2B4) and *Tnfrsf9* (4-1BB) (Fig. 1c,d and Extended Data Fig. 1c). The bifurcation of the two lineages occurred in the small, but distinct, T_transit_ cluster that is marked by the downregulation of Tpex-associated genes, including *Sell* (CD62L), *Ccr7*, *Tcf7*, *Il7r*, *Id3*, and *Slamf6* (Ly108), and an intermediate expression of terminal exhaustion markers (Fig. 1d). To further explore the metabolic-transcriptional programs during the differentiation of stem-like CD62L^+^ Tpex cells into CTL-like and terminally exhausted T cells, we calculated gene set enrichment scores for metabolic pathways, such as glycolysis, mitochondrial respiration, lipid metabolism and the pentose phosphate pathway (Supplementary Table S1). These analyses revealed a sharp decline in mitochondrial (gene set ID M9577) and respiratory chain gene expression (M19046) at the transition between Tpex to terminally differentiated T cells (Fig. 1e), indicating a relationship between mitochondrial (dys-) function and T cell exhaustion. In parallel to our scRNA sequencing data, we analyzed publicly available bulk RNA sequencing data of ID3^+^ Tpex and Tim-3^+^ Tex cells from chronically infected mice ^5^ (Extended Data Fig. 1d,e). Gene set enrichment analyses (GSEA) and clustering of gene networks revealed a significant correlation of pathways involved in *mitochondrial metabolism* and *translation* with Tpex cells, whereas the transcriptomes of Tex cells were primarily associated with *signal transduction, cell cycle* and *DNA repair* gene expression signatures (Extended Data Fig. 1d). The notion that Tpex cells rely more on mitochondrial respiration than Tex cells was further supported by GSEA using gene ontology (GO) pathways, such as *mitochondrial genes* (gene set ID M9577), *mitochondrial translation* (M27446) and *respiratory electron transport and ATP synthesis by chemiosmotic coupling* (M1025) (Extended Data Fig. 1e).

**Figure 1.**
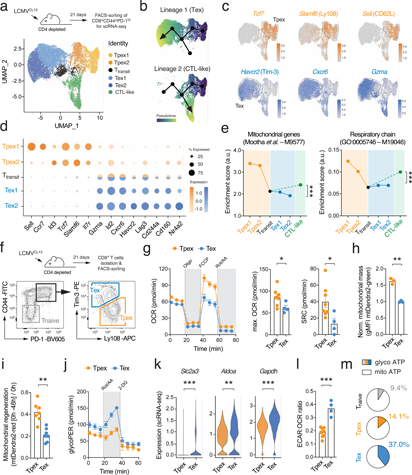
T cell exhaustion is characterized by metabolic reprogramming. **(a)** Naïve C57BL/6 mice were chronically infected with the LCMV strain clone 13 (LCMV^CL13^) and FACS-sorted CD8^+^CD44^+^PD-1^hi^ T cells were subjected to single-cell (sc) RNA sequencing 21 days post infection (d.p.i.). Uniform manifold approximation and projection (UMAP) visualization of ∼ 13.000 non-proliferating (*Mki67* negative) T cells coloured based on their classification into six clusters. **(b)** Prediction of developmental trajectories using slingshot analysis; cells are colour-coded according to pseudotime. **(c)** Normalised gene expression of *Tcf7*, *Slamf6* (Ly108), *Sell* (CD62L), *Havcr2* (Tim-3), *Cxcr6* and *Gzma* projected onto UMAP clusters. **(d)** Dot plot analysis of selected cluster markers representing Tpex and Tex cell subsets; color intensity and dot size represent z-score mean expression and percentage of cells expressing the gene, respectively. **(e)** Enrichment of mitochondrial and respiratory chain gene expression signatures in clusters representing Tpex, CTL-like and terminally exhausted (Tex) cells. **(f)** Flow cytometric analysis and cell sorting strategy of Tpex and Tex cells from chronically infected mice. **(g)** Analyses of oxygen consumption rate (OCR) and spare respiratory capacity (SRC) of Tpex and Tex cells using a Seahorse extracellular flux analyzer; means ± SEM of 4-7 experiments. **(h)** Analysis of mitochondrial content (mass/volume) in Tpex and Tex cells using LCMV^CL^^13^ infected mito-Dendra2 mice; means ± SEM of 3 mice. **(i)** Mitochondrial regeneration capacity of mito-Dendra2 Tpex and Tex cells after photoconversion *in vitro*; means ± SEM of 7 mice. **(j)** Glycolytic proton efflux rate (glycoPER) analyses of Tpex and Tex cells using a Seahorse extracellular flux analyzer; means ± SEM of 4-7 experiments. **(k)** Violin plots displaying *Slc2a3* (GLUT3), *Aldoa* and *Gapdh* gene expression in Tpex and Tex cells analyzed by scRNA-seq as shown in (A-E). **(l)** Ratio of extracellular acidification rate (ECAR) to oxygen consumption rate (OCR) in Tpex and Tex cells; means ± SEM of 4-7 experiments. **(m)** Relative contribution of glycolysis and mitochondrial respiration to cellular ATP production in naïve, Tpex and Tex cells in chronically infected mice; means ± SEM of 4-7 experiments. *, p<0.05; **, p<0.01; ***, p<0.001 by unpaired Student’s t-test in (g), (h), (i) and (l).

To directly measure mitochondrial respiration in exhausted T cells, we infected WT mice with LCMV^CL13^ and isolated Ly108^+^ Tpex and Tim-3^+^ Tex cells by FACS sorting (Fig. 1f). In line with the transcriptomic data, extracellular flux analyses revealed higher basal and maximal oxygen consumption rates (OCR) of Tpex compared to Tex cells (Fig. 1g), indicating that Tpex cells are more reliant on OXPHOS. The spare respiratory capacity (SRC), calculated as the difference between maximal and basal OCR, suggested that Tpex cells have a greater metabolic reserve to utilize mitochondrial respiration, whereas Tex cells operate OXPHOS almost at their maximal capacity (Fig. 1g). The difference in the SRC can be explained by an altered mitochondrial content and/or differences in their mitochondrial regenerative capacity ^31^. To test these possibilities, we employed mito-Dendra2 mice ^32^, which express a mitochondrial-localized version of the Dendra2 protein. Leveraging the green fluorescence of mito-Dendra2, which correlates with the mass/volume of the mitochondria ^32^, we could readily detect that the mitochondrial content of Tpex cells was significantly higher compared to Tex cells (Fig. 1h and Extended Data Fig. 2a). To delineate mitochondrial turnover and regeneration capacity in exhausted T cells, we labelled T cells of chronically infected mito-Dendra2 mice with cell trace violet (CTV) and irreversibly photo-switched their mitochondrial fluorescence emission spectrum from green to red fluorescence upon exposure to 405 nm laser light (Extended Data Fig. 2a,b). We tracked the emergence of non-red mitochondria in undivided Tpex and Tex cells *in vitro* as a measure for mitochondrial regeneration (Fig. 2i and Extended Data Fig. 2b). Mitotracker and TMRE validated the lower mitochondrial content and membrane potential in Tex compared to Tpex cells (Extended Data Fig. 2c). Furthermore, Tex cells also showed a diminished expression of several electron transport chain (ETC) complexes (Extended Data Fig. 2d,e) and mitochondrial biogenesis factors, such as PGC-1α and TFAM (Extended Data Fig. 2d), demonstrating that Tpex cells possess mitochondria with increased functionality and greater regenerative capacity compared to Tex cells.

**Figure 2.**
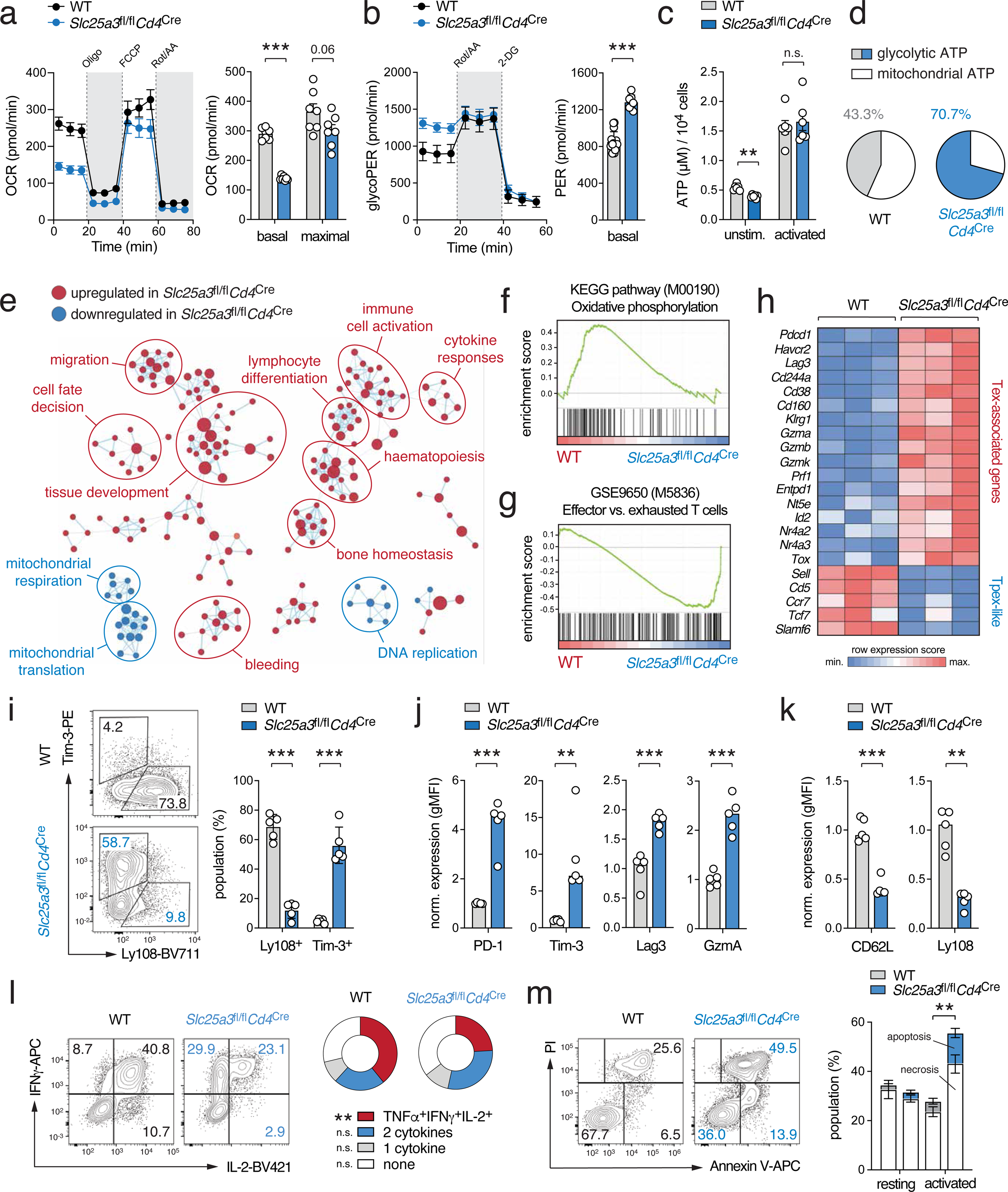
Genetic suppression of mitochondrial ATP production promotes glycolytic-transcriptional reprogramming of T cells. **(a, b)** Analyses of (a) oxygen consumption rate (OCR) and (b) glycolytic proton efflux rate (glycoPER) of activated WT and mPiC-deficient (*Slc25a3*^fl/fl^*Cd4*^Cre^) T cells at day 2 of culture using a Seahorse extracellular flux analyzer; means ± SEM of 7 mice. **(c)** Analysis of ATP concentrations in unstimulated and anti-CD3/CD28 activated WT and mPiC-deficient T cells; means ± SEM of 3 mice. **(d)** Relative contribution of glycolysis and mitochondrial respiration to cellular ATP production in anti-CD3/CD28 stimulated WT and mPiC-deficient T cells; means ± SEM of 3 mice. **(e)** Network clustering of RNA sequencing data of significantly (p < 0.05) enriched gene expression signatures between CTL-differentiated WT and mPiC-deficient T cells. Down- and upregulated gene signatures in mPiC-deficient T cells are shown in blue and red, respectively. **(f, g)** Gene set enrichment analysis (GSEA) of *oxidative phosphorylation* (KEGG pathway) (f) and *effector versus exhausted T cell* (GSE9650) gene signatures (g) in differentiated WT and mPiC-deficient T cells after 6 days of culture. **(h)** Heatmap expression analysis of selected genes in differentiated WT and mPiC-deficient CTLs after 6 days in culture. **(i-k)** Differentiation of WT and mPiC-deficient T cells into CTLs *in vitro*, means ± SEM of 5 mice. **(j, k)** Flow cytometric quantification of exhaustion (j) and memory (k) marker expression in WT and mPiC-deficient CTLs (day 6), means ± SEM of 5 mice. **(l)** Analysis of polyfunctional TNFα, IFNγ and IL-2 expression by WT and mPiC-deficient T cells after 6 days of culture and anti-CD3/CD28 restimulation; means ± SEM of 3 mice. **(m)** Analysis of apoptosis in resting and anti-CD3/CD28 stimulated T cells by flow cytometry; means ± SEM of 3 mice. *, p<0.05; **, p<0.01; ***, p<0.001 by unpaired Student’s t-test in (b), (c) and (i-m).

Intriguingly, and in contrast to their impaired mitochondrial performance, Tex cells showed an elevated glycolytic proton efflux rate (glycoPER) compared to Tpex cells (Fig. 1j), suggesting that Tex cells compensate their mitochondrial insufficiency by an elevated glycolytic activity. The increased extracellular acidification rate (ECAR) of Tex cells correlated with a greater expression of nutrient transporters and glycolytic enzymes, such as *Slc2a3* (encoding the glucose transporter GLUT3 ^33^), *Aldoa* and *Gapdh* (Fig. 1k). The notion that T cell exhaustion is mediated by a metabolic switch from mitochondrial respiration to aerobic glycolysis was further supported by a higher ECAR:OCR ratio in Tex cells compared to precursor cells (Fig. 1l). When we calculated the contribution of glycolytic and mitochondrial metabolism to ATP production, we found that Tpex cells rely primarily on OXPHOS, whereas Tex cells showed a ∼ 2.5-fold greater dependency on aerobic glycolysis (Fig. 1m).

Collectively, these data show that diminished mitochondrial respiration and a Warburg effect-like glycolytic reprogramming are metabolic hallmarks of terminal T cell exhaustion.

### Mitochondrial respiration controls the functional exhaustion of T cells

Having established that mitochondrial respiration is a hallmark of stem-like Tpex cells and that impaired mitochondrial ATP production coincides with the transition of Tpex into Tex cells, we set out to delineate if these metabolic changes are a consequence or the cause of T cell exhaustion. To directly address the role of mitochondrial bioenergetics, we generated mice with T cell-specific deletion of the *Slc25a3* gene that encodes for the mitochondrial phosphate carrier (mPiC) ^34^. Inactivation of mPiC causes a paucity of free inorganic phosphate within the mitochondrial matrix as a rate-limiting step in the biosynthesis of mitochondrial ATP (Extended Data Fig. 3a) ^34, 35^. Thus, *Slc25a3*^fl/fl^*Cd4*^Cre^ mice are an excellent genetic model to investigate the causality between mitochondrial ATP production and T cell exhaustion. Mice with T cell-specific mPiC ablation were born at the expected Mendelian ratio without obvious defects in the thymic αβ CD4^+^ and CD8^+^ T cell development (Extended Data Fig. 3b,c). However, *Slc25a3*^fl/fl^*Cd4*^Cre^ mice had significantly reduced conventional (Extended Data Fig. 3d) and regulatory T cells (Extended Data Fig. 3e) in their peripheral lymphoid organs. The phenotype of the CD4^+^ and CD8^+^ T cells in *Slc25a3*^fl/fl^*Cd4*^Cre^ mice was slightly shifted towards CD44^+^CD62L^−^ effector T cells at the expense of naïve and CD44^+^CD62L^+^ central memory cells (Extended data Fig. 3f,g). To examine the metabolic profiles of mPiC-deficient T cells, we stimulated naive CD8^+^ T cells of with anti-CD3/CD28 for 2 days (activation) followed by an incubation with IL-2 for additional 4 days (CTL differentiation) (Fig. 2a-d and Extended Data Fig. 2h,i). Seahorse extracellular flux analyses revealed markedly reduced basal and maximal OCR levels in anti-CD3/CD28 activated mPiC-deficient T cells (Fig. 2a) coinciding with higher glycolytic activity (Fig. 2b). This altered metabolic profile was also observed in fully differentiated CTLs (Extended Data Fig. 3h,i). Consistent with impaired mitochondrial respiration in mPiC-deficient T cells, we also observed lower glucose-derived ^13^C-labelling of TCA cycle intermediates, whereas the fractional enrichment of ^13^C in glycolytic metabolites was unaltered or increased (Extended Data Fig. 3j). We detected significantly higher ^13^C-lactate levels, confirming a ‘Warburg effect-like’ shifted glucose utilization in mPiC-deficient T cells (Extended Data Fig. 3k). Cellular ATP levels were unaltered in mPiC-deficient T cells upon antigen receptor stimulation (Fig. 2c), suggesting that elevated aerobic glycolysis compensates for the impaired mitochondrial ATP biosynthesis. This notion is in line with a higher ECAR:OCR ratio (Extended Data Fig. 3i) and a greater bioenergetic dependency on aerobic glycolysis of mPiC-deficient T cells (Fig. 2d). The observation that both Tex (*ex vivo*) and mPiC-deficient T cells (*in vitro*) prefer aerobic glycolysis over mitochondrial respiration highlights the potential of *Slc25a3*^fl/fl^*Cd4*^Cre^ mice to explore the mechanistic link between metabolic reprogramming and T cell exhaustion.

To further investigate this hypothesis, we performed bulk RNA sequencing of activated (day 2) and CTL-differentiated T cells (day 6) to delineate how metabolic reprogramming affects gene expression of mPiC-deficient T cells (Extended Data Fig. 3l,m). The gene expression profiles of both genotypes were clearly distinct and revealed 1475 and 463 differentially expressed genes (DEGs) in activated T cells and CTLs, respectively (Extended Data Fig. 3l,m). To explore the physiological processes governed by mPiC-dependent metabolic reprogramming, we performed pathway enrichment and network cluster analyses (Fig. 2e). Surprisingly, gene signatures correlating with *immune cell activation* and *lymphocyte differentiation* were enriched in mPiC-deficient T cells after activation (Fig. 2e), suggesting that T cell activation is not impaired, but rather enhanced, when mitochondrial ATP synthesis is substituted by aerobic glycolysis. Indeed, gene expression signatures involved in *mitochondrial respiration*, *mitochondrial translation* and *oxidative phosphorylation* were lower in mPiC-deficient T cells (Fig. 2e,f). The gene expression profiles of mPiC-deficient T cells were poorly correlated with those of effector T cells but were shifted towards an exhaustion signature (Fig. 2g), indicating that metabolic reprogramming promotes the terminal differentiation of T cells. Numerous genes that are characteristic for T cell exhaustion, such as *Havcr2*, *Pdcd1*, *Lag3* and *Gzma*, were markedly upregulated in mPiC-deficient CTLs (day 6), whereas Tpex-like genes, including *Sell*, *Ccr7*, *Tcf7* and *Slamf6*, were downregulated (Fig. 2h). Consistent with the transcriptomic findings, Tim-3, PD-1, Lag3 and granzyme A protein levels were also higher in mPiC-deficient CTLs compared to WT controls (Fig. 2i,j). Intriguingly, when we differentiated WT and mPiC-deficient T cells with IL-7 plus IL-15 to generate ‘memory-like’ T cells *in vitro*, we did not observe an elevated expression of co-inhibitory receptors (Extended Data Fig. 4a-c). Nonetheless, CD62L and Ly108 expression were still impaired in mPiC-deficient memory-like T cells (Extended Data Fig. 4b), indicating that mitochondrial respiration supports the maintenance of both Tpex and memory-like T cells. In addition to the ‘acute’ activation of T cells with anti-CD3/CD28 followed by resting in IL-2 or IL-15 (Fig. 2i, j and Extended Data Fig. 4a-c), we also stimulated WT and mPiC-deficient T cells ‘chronically’ to provoke exhaustion by continuous antigenic stimulation ^24^ (Extended Data Fig. 4d-g). As expected, chronic stimulation massively upregulated the expression of activation markers and co-inhibitory receptors (Extended Data Fig. 4f). Although many of these markers were already maximally expressed in under these conditions, Tim-3 expression was significantly higher in mPiC-deficient T cells compared to WT controls (Extended Data Fig. 4f), whereas their expression of CD62L and Ly108 was impaired (Extended Data Fig. 4e). Of note, the upregulation of co-inhibitory receptors alone does not prove functional exhaustion. To demonstrate that mPiC-deficient T cells are indeed dysfunctional, we analyzed the cytokine production profile (Fig 2l and Extended Data Fig. 2g) and apoptosis (Fig. 2m). As expected, mPiC-deficient T cells produced significantly less IFNγ and IL-2 under ‘acute’ and ‘chronic’ culture conditions (Fig 2l and Extended Data Fig. 4g) and showed impaired survival after antigen receptor ligation (Fig. 2m). Collectively, these data demonstrate that mPiC ablation causes both phenotypic and functional exhaustion of T cells.

Based on the observation that mitochondrial (dys-) function affects the expression of both stemness and exhaustion markers *in vitro* (Fig. 2h-k), we next investigated how mitochondrial insufficiency affects T cell differentiation during chronic viral infection *in vivo*. To track antigen-specific T cells, we crossed WT and *Slc25a3*^fl/fl^*Cd4*^Cre^ mice to P14 animals that express an LCMV-specific T cell receptor ^36^ and fluorescent reporters. 14 days after adoptive co-transfer of WT (GFP^+^) and mPiC-deficient P14 T cells (tdTomato^+^) into chronically infected WT host mice, we analyzed the proportions of Ly108^+^ Tpex and Tim-3^+^ Tex cells among the donor T cell populations (Fig. 3a). The frequencies of Tpex cells were ∼ 50% reduced in mPiC-deficient T cells compared to those of WT T cells, whereas the proportion of Tex cells was reciprocally increased in absence of mPiC (Fig. 3a and Extended Data Fig. 5a). Of note, GFP^+^ WT T cells numerically outcompeted tdTomato^+^ mPiC-deficient P14 cells over time and 21 days after infection we could barely detect mPiC-deficient T cells in the spleens and LNs of the recipient mice (Extended Data Fig. 5b,c). The competitive disadvantage of mPiC-deficient T cells may be explained by an impaired clonal expansion, elevated apoptosis or both. Indeed, mPiC-deficient T cells showed both impaired proliferation (Extended Data Fig. 5d) and increased apoptosis (Fig. 2m), which also affected their clonal expansion *in vitro* (Extended Data Fig. 5e). Adoptive co-transfer experiments of cell trace violet (CTV)-labelled P14 T cells confirmed a reduced proliferative capacity of mPiC-deficient T cells *in vivo* (Extended Data Fig. 5f), providing a plausible explanation for their competitive disadvantage. These findings suggest that mitochondrial respiration is required for the maintenance of Tpex cells during chronic infection and/or that mitochondrial insufficiency accelerates terminal T cell exhaustion. To test the latter hypothesis, we infected WT and *Slc25a3*^fl/fl^*Cd4*^Cre^ mice with the acute ‘Armstrong’ strain of LCMV (LCMV^ARM^), which provokes a transient infection in WT mice without causing T cell exhaustion ^37^. T cells of LCMV^ARM^ infected WT mice displayed the expected effector phenotype and the few PD-1^+^ T cells co-expressed Ly108 (Fig. 3b). By contrast, the frequency of PD-1^+^ T cells was dramatically increased in *Slc25a3*^fl/fl^*Cd4*^Cre^ mice, of which ∼ 60% were positive for Tim-3 (Fig. 3b). We also tested the functional capacities of WT and mPiC-deficient T cells after LCMV infection by determining their cytokine profiles. mPiC-deficient T cells showed a markedly reduced capability to produce two or more cytokines after PMA/iono restimulation (Fig. 3c,d), demonstrating that defective mitochondrial respiration not only causes phenotypic features of T cell exhaustion but also results in their functional impairment. To test the hypothesis that augmented mitochondrial respiration conversely sustains the stemness of exhausted T cells, we retrovirally (RV) overexpressed mPiC in naïve T cells and adoptively co-transferred control (pMIG-Ametrine^+^) and mPIC-overexpressing P14 T cells (pMIG-SLC25A3-GFP^+^) into LCMV^CL^^13^ infected recipient mice (Fig. 3e). Although both donor T cell populations expanded comparably in the 7 days post infection, mPiC-overexpressing T cells outcompeted control cells over time (Fig. 3f-h), suggesting that augmented mitochondrial respiration (Fig. 3i-k) protects virus-specific T cells from functional exhaustion. Indeed, phenotypic and functional analyses of donor T cells revealed that mPiC-overexpressing T cells retained a higher proportion of Tpex cells (Fig. 3l) and a superior capacity to produce cytokines (Fig. 3m).

**Figure 3.**
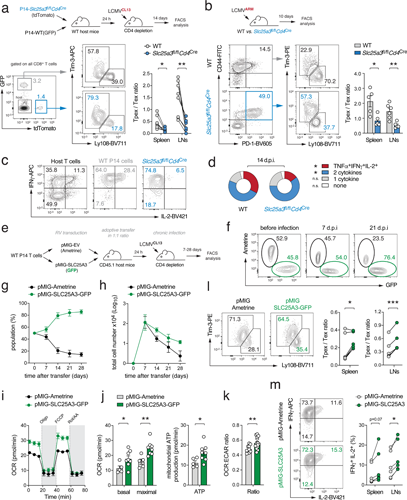
Mitochondrial respiration controls the functional exhaustion of virus-specific T cells. **(a)** Adoptive co-transfer of naïve GFP^+^ WT and tdTomato^+^ mPiC-deficient (*Slc25a3*^fl/fl^*Cd4*^Cre^) P14 T cells into C57BL/6 mice before chronic infection with LCMV clone 13 (LCMV^CL13^). Flow cytometric analysis of Tpex and Tex cells in spleen and LNs of the host mice 14 days post infection (d.p.i.); n = 8 mice. **(b)** Acute infection of WT and *Slc25a3*^fl/fl^*Cd4*^Cre^ mice with LCMV Armstrong (LCMV^ARM^). Analysis of Tpex and Tex cells in spleen and LNs was performed 10 d.p.i.; means ± SEM of 5 mice. **(c, d)** Analysis of polyfunctional TNFα, IFNγ and IL-2 expression after PMA/iono restimulation in T cells of WT and mPiC-deficient P14 T cells after co-transfer into chronically infected mice; means ± SEM of 6-10 mice. **(e-m)** Ectopic expression of mPiC attenuates T cell exhaustion. **(e)** Retroviral transduction of WT P14 T cells with SLC25A3/mPiC (GFP^+^) or pMIG empty vectors (Ametrine^+^) followed by adoptive co-transfer into chronically infected mice. **(f)** Representative flow cytometric analysis of GFP^+^ and Ametrine^+^ donor P14 T cells after transfer into LCMV^CL^^13^ infected CD45.1^+^ mice. **(g, h)** Relative (g) and absolute (h) numbers of GFP^+^ and Ametrine^+^ donor P14 T cells in the spleens of recipient mice; means ± SEM of 2-6 mice per timepoint. **(i, j)** Oxygen consumption rate (OCR) (i) and mitochondrial ATP production (j) in mPiC (GFP^+^) and empty vector (Ametrine^+^) transduced P14 cells *ex vivo* 7 d.p.i. using a Seahorse extracellular flux analyzer; means ± SEM of 3 biological samples in technical replicates. **(k)** Ratio of OCR to extracellular acidification rate (ECAR) in mPiC overexpressing *versus* empty vector T cells; means ± SEM of 3 biological samples in technical replicates. **(l)** Flow cytometric analysis of Tpex and Tex cell ratio in mPiC (GFP^+^) and empty vector transduced (Ametrine^+^) P14 cells in the spleen and LNs of the host mice 7 d.p.i.; n = 6 mice. **(m)** IFNγ and IL-2 expression in mPiC (GFP^+^) and empty vector transduced (Ametrine^+^) P14 cells after PMA/iono restimulation 7 d.p.i.; means ± SEM of 6 mice. *, p<0.05; **, p<0.01; ***, p<0.001 by paired and unpaired Student’s t-test in (a), (b), (d) and (i-m), respectively.

Taken together, these findings demonstrate that defects in mitochondrial respiration are sufficient to elicit the transcriptional, phenotypic and functional features of T cell exhaustion even in the absence of continuous antigen exposure.

### Oxidative stress stabilizes HIF-1α protein levels in Tpex cells

To understand how metabolism controls the complex transcriptional processes during T cell exhaustion, we profiled the metabolome of *in vitro* activated WT and mPiC-deficient T cells by untargeted liquid chromatography and mass spectrometry (LC/MS) (Fig. 4a). Among the most downregulated metabolites in mPiC-deficient T cells were several glycolytic intermediates, such as *fructose-6-phosphate*, *fructose-1,6-bisphospate*, *dihydroxyacetone-phosphate* (DHAP), *glyceraldehyde-3-phosphate* (G3P) and *phosphoenolpyruvate* (PEP), supposably due to an accelerated glycolytic flux. In addition, the reduced form of *nicotinamide adenine dinucleotide phosphate* (NADPH) was downregulated in mPiC-deficient T cells (Fig. 4a), resulting in a shifted NADPH/NADP^+^ ratio (Fig. 4b). The regeneration of cytosolic NADPH is linked to the pentose phosphate pathway (PPP) and metabolically coupled to the conversion of glucose-6-phosphate (G6P) to pentose-5-phosphate ^38^. We next used stable-isotope tracing to determine the glucose-derived ^13^C-labelling pattern of TCA cycle and PPP intermediates. Intriguingly, ^13^C-pentose-5-phosphate was markedly reduced in mPiC-deficient T cells (Extended Data Fig. 6a) and the gene expression signature of the PPP declined progressively during T cell exhaustion (Extended Data Fig. 6b). These findings indicate that glucose is re-directed into glycolysis to compensate for the impaired mitochondrial ATP production rate, thereby limiting NADPH production. Given the crucial role of NADPH for the cellular redox hemostasis as an antioxidant ^38^, we hypothesized that oxidative stress promotes the exhaustion phenotype of mPiC-deficient T cells. This notion was supported by GSEA that revealed a strong correlation with the *reactive oxygen species (ROS) pathway* gene expression signature (gene set ID M5938) (Fig. 4c) and an upregulation of genes involved in ROS detoxification (Fig. 4d). Flow cytometric analyses using CellROX and MitoSox revealed elevated cytosolic and mitochondrial ROS levels, respectively (Fig. 4e). To test whether neutralization of cellular ROS can rescue mPiC-deficient T cells from functional exhaustion, we supplemented WT and mPiC-deficient T cells with N-acetylcysteine (NAC), a cell-permeant form of cysteine that prevents redox stress via the biosynthesis of glutathione ^39^. In line with recent reports ^23, 24^, NAC normalized cellular ROS levels in mPiC-deficient T cells and prevented the expression of exhaustion markers (Fig. 4f). NAC also restored their capacity to produce cytokines (Fig. 4g). These data demonstrate that cellular redox regulation is a critical molecular rheostat linking metabolic reprogramming to T cell exhaustion.

**Figure 4.**
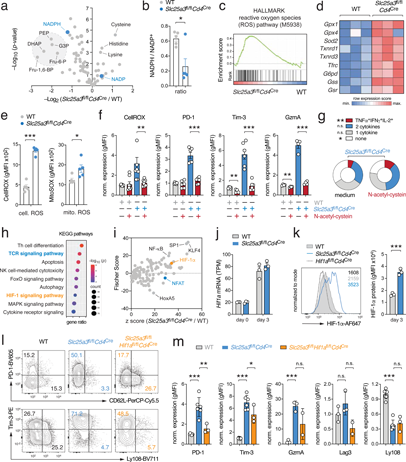
Mitochondrial insufficiency causes ROS-mediated HIF-1α protein stabilization. **(a)** Volcano plot of differential metabolite concentrations of *in vitro* activated WT and mPiC-deficient (*Slc25a3*^fl/fl^*Cd4*^Cre^) T cells analyzed by untargeted liquid chromatography and mass spectrometry (LC/MS); 4 biological replicates per group. **(b)** Analysis of NADPH/NADP^+^ ratios in WT and mPiC-deficient T cells using LC/MS; 4 biological replicates per group. **(c)** Gene set enrichment analysis (GSEA) of the *reactive oxygen species (ROS) pathway* gene signature in WT and mPiC-deficient T cells. **(d)** Heatmap analysis of selected genes involved in ROS detoxification. **(e)** Flow cytometric analysis of cellular (CellROX) and mitochondrial ROS (MitoSox) in WT and mPiC-deficient T cells; means ± SEM of 5 mice. **(f)** Scavenging of ROS by N-acetylcysteine (NAC) prevents exhaustion of mPiC-deficient T cells. Flow cytometric analyses of CellRox, PD-1, Tim-3 and granzyme A expression in WT and mPiC-deficient T cells treated with NAC; means ± SEM of 4-8 mice. **(g)** Analysis of polyfunctional TNFα, IFNγ and IL-2 expression after PMA/iono restimulation of mPiC-deficient T cells treated with or without NAC; means ± SEM of 5 mice. **(h)** Kyoto Encyclopedia of Genes and Genomes (KEGG) pathway enrichment analysis using bulk RNA sequencing data of WT and mPiC-deficient T cells. **(i)** Upstream transcription factor prediction analysis using differentially expressed genes (DEGs) between WT and mPiC-deficient T cells. **(j)** Analysis of *Hif1a* gene expression in WT and mPiC-deficient T cells by RNA sequencing; 3 biological replicates per group. **(k)** Flow cytometric analysis of HIF-1α protein expression in WT, mPiC- and HIF-1α-deficient T cells; means ± SEM of 3 mice. **(l, m)** Differentiation of WT, mPiC-deficient (*Slc25a3*^fl/fl^*Cd4*^Cre^) and mPiC/HIF-1α double-deficient (*Slc25a3*^fl/fl^*Hif1a*^fl/fl^*Cd4*^Cre^) T cells *in vitro*. Representative flow cytometric analysis (l) and quantification of PD-1, Tim-3, granzyme A, Lag3 and Ly108 expression on WT, mPiC-deficient and mPiC/HIF-1α double-deficient T cells (m); means ± SEM of 2-7 mice. *, p<0.05; **, p<0.01; ***, p<0.001 by unpaired Student’s t-test in (b), (e), (f), (g), (k) and (m).

We next addressed how elevated ROS levels are ‘translated’ into gene expression in mPiC-deficient T cells. To identify potential signaling pathways and transcriptional regulators, we performed pathway (Fig. 4h) and transcription factor enrichment analyses (Fig. 4i). Both analyses highlighted an involvement of TCR signaling and NFAT (Fig. 4h,i). This was intriguing because NFAT controls numerous genes involved in T cell exhaustion ^18, 40, 41, 42^. It has been shown that mitochondrial-derived ROS can sustain the nuclear translocation of NFAT during T cell activation and promote exhaustion ^23, 24, 43, 44^ (Extended Data Fig. 6c). Chronically stimulated T cells express predominantly the short isoform of NFATc1 ^45^, involved in lymphocyte activation, differentiation and effector function ^18, 46^. To test whether NFATc1 also drives exhaustion of mPiC-deficient CTLs, we crossed *Nfatc1*^fl/fl^ to *Slc25a3*^fl/fl^*Cd4*^Cre^ mice to obtain mPiC/NFATc1 double-deficient T cells. Surprisingly, ablation of NFATc1 did not prevent the upregulation of PD-1, Tim-3 and other co-inhibitory receptors in mPiC-deficient T cells (Extended Data Fig. 6d,e), suggesting that different NFAT family members compensate for the loss of NFATc1 and/or that other transcription factors promote exhaustion of mPiC-deficient T cells. Supporting the latter, pathway and transcription factor enrichment analyses identified HIF-1α as a potential regulator involved (Fig. 4h,i). Gene and protein levels of HIF-1α were markedly increased in Tpex and transitory T cells (Extended Data Fig. 6f,g). Furthermore, HIF-1α target gene expression peaked during the transition of Tpex into Tex cells (Extended Data Fig. 6h), indicating that HIF-1α-mediated gene expression contributes to T cell exhaustion. The activity of HIF-1α is regulated at the transcriptional and posttranslational level ^47, 48^. Protein stability of HIF-1α is dependent on cellular oxygen tension because prolyl hydoxylases (PHDs) catalyse the hydroxylation of HIF-1α in normoxia, which promotes its ubiquitination and proteasomal degradation. Under hypoxia – or when the enzymatic activity of PHD is inhibited – HIF-1α protein is stabilized and dimerizes with ARNT to form an active transcription factor complex ^47, 48^. Importantly, PHDs require α-ketoglutarate, Fe^2+^ and ascorbate as cofactors and ROS-mediated oxidation of these cofactors inhibits the enzymatic activity of PHD (Extended Data Fig. 6i) ^48, 49^. While gene expression of *Hif1a* was comparable between WT and mPiC-deficient T cells *in vitro* (Fig. 4j), HIF-1α protein levels were dramatically increased in mPiC-deficient T cells (Fig. 4k). The hydroxylation of HIF-1α is metabolically coupled to the conversion of α-ketoglutarate to succinate, which was reduced in mPiC-deficient T cells (Extended Data Fig. 6j). Importantly, scavenging ROS by NAC normalized HIF-1α protein expression levels in mPiC-deficient T cells (Extended Data Fig. 6k), supporting the notion that ROS levels prevent the proteasomal degradation of HIF-1α protein. To examine whether HIF-1α promotes to the upregulation of exhaustion-associated proteins in mPiC-deficient T cells, we crossed *Hif1a*^fl/fl^ to *Slc25a3*^fl/fl^*Cd4*^Cre^ mice to generate mPiC/HIF-1α double-deficient T cells. Ablation of HIF-1α attenuated the expression of PD-1, Tim-3 and Lag3 in mPiC-deficient T cells, but failed to restore the expression of Ly108 (Fig. 4l,m).

In addition to mPiC-deficient mice, we established an additional genetic model of mitochondrial dysfunction by generating mice with T cell-specific ablation of TFAM (*Transcription factor A, mitochondrial*) (Extended Data Fig. 7). TFAM was highly expressed in Tpex cells but almost completely lost in terminally exhausted T cells (Extended Data Fig. 2d). Because TFAM plays a crucial role in regulating mitochondrial DNA transcription and replication, its ablation efficiently inhibits mitochondrial biogenesis and oxidative phosphorylation ^50, 51^. Similar as observed after mPiC inactivation, TFAM-deficient T cells also showed the characteristic metabolic switch from mitochondrial respiration to aerobic glycolysis (Extended Data Fig. 7a,b), accumulation of cellular ROS (Extended Data Fig. 7c) and the posttranslational stabilization of HIF-1α (Extended Data Fig. 7d), which was correlated with an upregulation of different exhaustion markers (Extended Data Fig. 7e,f).

Collectively, these data demonstrate that oxidative stress and metabolic reprogramming due to mitochondrial insufficiency supports HIF-1α protein stabilization and T cell exhaustion.

### HIF-1α-mediated glycolytic reprogramming drives terminal T cell differentiation

We next investigated how HIF-1α-mediated gene expression affects terminal T cell exhaustion during chronic viral infection. We crossed *Hif1a*^fl/fl^*Cd4*^Cre^ mice to P14 animals and adoptively co-transferred equal numbers of WT (GFP^+^) and HIF-1α-deficient T cells (tdTomato^+^) into chronically infected WT host mice. 21 days post infection, we retrieved GFP^+^ and tdTomato^+^ donor T cells and subjected individually barcoded samples to multiplexed scRNA sequencing analysis (Extended Data Fig. 8a). Using a similar analysis strategy as for polyclonal T cells (Fig. 1a-e), we identified UMAP clusters representing precursor and terminally exhausted T cells (Fig. 5a,b and Extended Data Fig. 8b,c). Both precursor clusters expressed the ‘stemness’ markers *Tcf7* and *Slamf6*, but Tpex2 differed from Tpex1 cells by the loss of *Sell* and *Il7r* (Fig. 5b,c and Extended Data Fig. 8b,c). The two terminally exhausted T cell populations were characterized by the expression of *Havcr2*, *Cxcr6*, *Id2*, *Lag3*, *Gzma* and *Cd244a* (Fig. 5b,c and Extended Data Fig. 8b,c). Diffusion pseudotime analysis predicted a unidirectional trajectory originating from stem-like CD62L^+^ Tpex1 cells ^10^ into terminally differentiated Tex cells (Fig. 5a). Consistent with the findings in polyclonal T cells (Fig. 1e), P14 T cells also showed a sharp decline in their mitochondrial gene signatures when progressing from Tpex to Tex cells (Extended Data Fig. 8d). Importantly, HIF-1α-deficient T cells were enriched within the Tpex populations, whereas most WT T cells were found in terminally exhausted UMAP clusters (Fig. 5d). Comparing the DEGs between WT and HIF-1α-deficient P14 T cells across clusters, we found that *Tcf7* and *Slamf6* were significantly higher in HIF-1α-deficient T cells (Fig. 5e), whereas exhaustion-associated genes were decreased compared to WT T cells (Fig. 5e). To test the hypothesis that HIF-1α-mediated gene expression supports the transition of Tpex into terminally exhausted T cells, we adoptively co-transferred WT and HIF-1α-deficient P14 T cells into chronically infected WT host mice. 14 days after transfer, we found a significantly increased Ly108^+^ Tpex cell population, but fewer terminally exhausted T cells, among HIF-1α-deficient donor T cells compared to WT controls (Fig. 5f). Similarly, when we analyzed endogenous T cell responses in WT and *Hif1a*^fl/fl^*Cd4*^Cre^ mice after chronic LCMV^CL^^13^ infection, we also observed a higher Tpex-to-Tex ratio among the virus-specific (GP_33-41_ tetramer^+^) T cells in HIF-1α-deficient mice (Extended Data Fig. 8e,f). These data suggest that HIF-1α-mediated gene expression contributes to the conversion of Tpex into terminally exhausted T cells.

**Figure 5.**
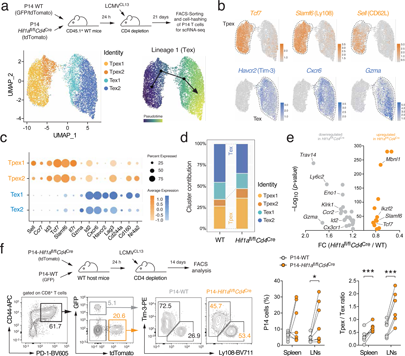
HIF-1α controls terminal differentiation of virus-specific T cells. **(a)** Adoptive co-transfer of naïve tdTomato^+^ GFP^+^ WT and tdTomato^+^ HIF-1α-deficient (*Hif1a*^fl/fl^*Cd4*^Cre^) P14 T cells into C57BL/6 mice before chronic infection with LCMV clone 13 (LCMV^CL13^). 21 days post infection (d.p.i.), donor P14 WT and HIF-1α-deficient T cells were FACS sorted, barcoded and multiplexed in a 1:1 ratio and and subjected to single-cell (sc) RNA sequencing. Uniform manifold approximation and projection (UMAP) visualization of ∼ 8.400 non-proliferating (*Mki67* low/negative) T cells identified four clusters. Prediction of developmental trajectories using slingshot analysis; colour-coding according to pseudotime. **(b)** Normalised gene expression of *Tcf7*, *Slamf6* (Ly108), *Sell* (CD62L), *Havcr2* (Tim-3), *Cxcr6* and *Gzma* projected onto UMAP clusters. **(c)** Dot plot analysis of selected cluster markers representing Tpex and Tex cell subsets; color intensity and dot size represent z-score mean expression and percentage of cells expressing the gene, respectively. **(d)** Relative contribution of individual UMAP clusters in WT and Hif-1α-deficient P14 T cells. **(e)** Volcano plot of differentially expressed genes (DEGs) between WT and HIF-1α-deficient P14 T cells using scRNA sequencing. **(f)** Adoptive co-transfer of naïve GFP^+^ WT and tdTomato^+^ HIF-1α-deficient P14 T cells into C57BL/6 mice before chronic infection with LCMV^CL13^. Flow cytometric analysis of Tpex and Tex cells in spleen and LNs of the host mice was performed 14 d.p.i.; n = 5-12 mice. *, p<0.05; ***, p<0.001 by paired Student’s t-test in (f).

To decipher how HIF-1α controls exhaustion of virus-specific T cells, we performed pathway enrichment analyses based on the DEGs between WT and HIF-1α-deficient P14 T cells (Fig. 5a-e). Besides the expected association with the *HIF-1α signaling pathway*, we found a strong correlation of the DEGs with the *glycolysis and gluconeogenesis* gene signature (Fig. 6a). The expression of rate-limiting enzymes of the glycolytic pathway, such as *phosphofructo kinase* (*Pfkl*), *aldolase* (*Aldoa*), *glyceraldehyde-3-phosphate dehydrogenase* (*Gapdh*), *enolase* (*Eno1*) and *pyruvate kinase* (*Pkm*), were significantly downregulated in HIF-1a-deficient Tpex and Tex cells (Fig. 6b,c). HIF-1α-deficient T cells also showed a ∼30% reduced glycolytic activity under hypoxic culture conditions, which was correlated with a lower expression of co-inhibitory receptors (Fig. 6d,e and Extended Data Fig. 9a,b). Intriguingly, HIF-1α-deficient CTLs compensated their glycolytic deficit by OXPHOS (Fig. 6f), as previously described for regulatory T cells ^52^. The higher bioenergetic dependency on mitochondrial respiration (Fig. 6g) correlated with an increased expression of CD62L and Ly108 (Extended Data Fig. 9b). This suggests that HIF-1α-mediated glycolytic gene expression and the ‘Warburg effect-like’ metabolic reprogramming promotes the differentiation from Tpex into Tex cells. Indeed, 2-desxoxyglucose (2-DG), a potent inhibitor of glycolysis, prevented the upregulation of Tim-3, Lag-3 and granzyme A expression in mPiC-deficient T cells (Extended Data Fig. 9c). To test whether glycolytic restriction also preserves the stemness of virus-specific T cells, we stimulated P14 T cells with anti-CD3/CD28 *in vitro* followed by 2-DG treatment for 24 h before we co-transferred equal numbers of 2-DG-treated (expressing tdTomato^+^ and GFP^+^) and control (GFP^+^) T cells into LCMV^CL^^13^ infected host mice (Fig. 6h and Extended Data Fig. 9d-f). 2-DG-treated P14 cells not only outcompeted control T cells 10 and 21 days after transfer (Fig. 6h and Extended Data Fig. 9f) but also retained more Tpex cells at both timepoints (Fig. 6h and Extended Data Fig. 9d,e). Of note, 2-DG-treated T cells were fully functional after transfer into chronically infected mice (Fig. 6i and Extended Data Fig. 9g), demonstrating that short-term inhibition of glycolysis preserves the stemness of virus-specific T cells without compromising their effector functions. To further address whether 2-DG treatment also augments the antitumor immunity of T cells, we stimulated ovalbumin (OVA)-specific OT-1 T cells with anti-CD3/CD28 for 2 days in vitro followed by 2-DG treatment for 24 h before transfer into mice bearing MC38 colon adenocarcinomas expressing OVA (Extended Data Fig. 9h-l). Short term-treatment of OT-1 T cells with 1 mM 2-DG increased the OCR:ECAR ratio (Extended Data Fig. 9i) and their bioenergetic dependency on mitochondrial respiration (Extended Data Fig. 9j), which correlated with a lower expression of co-inhibitory receptors (Extended Data Fig. 9k) and a greater antitumor efficacy (Extended Data Fig. 9l). These findings provide a rationale for the generation of ‘exhaustion-resistant’ CAR T cells for cancer immunotherapy. To test this concept in a murine model, we transduced naive T cells with a second-generation CAR targeting human CD19 ^42^. After retroviral transduction with the CAR construct, activated T cells were treated with 2-DG for 24 h before a suboptimal number of 1×10^6^ CAR T cells was adoptively transferred into lymphopenic host mice bearing hCD19-expressing MC38 tumors (Fig. 6j). Similar as observed for OVA-specific CTLs, 2-DG pretreated CAR T cells also showed an increased antitumor activity, resulting in improved tumor regression (Fig. 6k) and prolonged survival (Fig. 6l).

**Figure 6.**
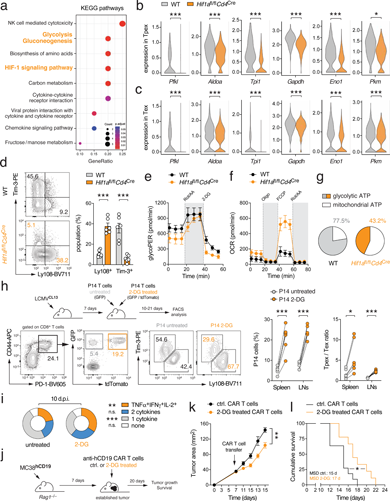
HIF-1α-mediated glycolytic reprogramming promotes T cell exhaustion. **(a)** Kyoto Encyclopedia of Genes and Genomes (KEGG) pathway enrichment analysis using single cell (sc) RNA sequencing of WT and HIF-1α-deficient T cells. **(b, c)** Violin plots displaying *Pfkl, Aldoa, Tpi1, Gapdh*, *Eno1* and *Pkm* gene expression in Tpex (b) and Tex cells (c) of WT and HIF-1α-deficient (*Hif1a*^fl/fl^*Cd4*^Cre^) P14 T cells analyzed by scRNA sequencing. **(d)** Differentiation of WT and HIF-1α-deficient T cells under hypoxia *in vitro*, means ± SEM of 6 mice. **(e, f)** Analyses of glycolytic proton efflux rate (glycoPER) (e) and oxygen consumption rate (OCR) (f) in WT and HIF-1α-deficient T cells using a Seahorse extracellular flux analyzer; means ± SEM of 3 mice. **(g)** Relative contribution of glycolysis and mitochondrial respiration to cellular ATP production in WT and HIF-1α-deficient T cells; means ± SEM of 3 mice. **(h, i)** Inhibition of glycolysis sustains the stemness of virus-specific T cells. **(h)** Adoptive co-transfer of 2-DG pre-treated (tdTomato^+^ GFP^+^) and control (GFP^+^) P14 T cells into C57BL/6 mice after chronic infection with LCMV^CL^^13^. Flow cytometric analysis of Tpex and Tex cells among donor P14 cells in spleen and LNs 10 d.p.i.; n = 6 mice. **(i)** Analysis of polyfunctional TNFα, IFNγ and IL-2 expression after PMA/iono restimulation of 2-DG-treated and control P14 T cells 10 days after co-transfer into chronically infected mice; means ± SEM of 6 mice. **(j-l)** 2-DG treatment during manufacturing augments the antitumor efficacy of CAR T cells. **(j)** 1×10^6^ anti-hCD19 CAR T cells treated with or without 2-DG for 24 h were adoptively transferred into *Rag1*^−/–^ mice 7 days after MC38^hCD19^ tumor inoculation. **(k, l)** Analysis of tumor growth (k) and cumulative survival (l) of MC38^hCD19^-bearing host mice after transfer of 2-DG or control-treated anti-hCD19 CAR T cells; means ± SEM of 9-11 mice. *, p<0.05; **, p<0.01; ***, p<0.001 by unpaired and paired Student’s t-test in (b-d) and (h). *, p<0.05 and **, p<0.05 in (k) and (l) by 2-way ANOVA and Mantel-Cox test, respectively.

All together, these findings suggest that both mitochondrial insufficiency and HIF-1α-mediated glycolytic reprogramming contribute to T cell exhaustion and that pharmacological inhibition of glycolysis is a promising metabolic intervention strategy to maintain stemness, longevity and functionality of (CAR) T cells during chronic viral infection and cancer immunotherapy.

## DISCUSSION

Exhausted T cells are heterogenous and comprise developmentally and functionally distinct subsets that differ in their transcriptional, epigenetic and metabolic signatures ^10, 23, 24, 25, 26, 27, 28, 30, 53^. By analyzing single-cell gene expression profiles of exhausted T cell subsets from chronically infected mice, we found that the transition of stem-like Tpex into functionally exhausted T cells is paralleled by a sharp decline in mitochondrial gene expression and a metabolic reprogramming towards aerobic glycolysis. Using novel genetic mouse models of mitochondrial insufficiency in T cells, we here show that this metabolic switch is sufficient to induce the exhaustion-associated gene expression program in Tpex cells, thus resembling an important cell-intrinsic trigger and not merely the consequence of exhaustion. In fact, mitochondrial insufficiency elicited T cell exhaustion even in the absence of continued antigen exposure. Mitochondrial decline and the re-direction of glucose into aerobic glycolysis diminishes the generation of NADPH, causing redox stress and an accumulation of ROS. Pathway and transcription factor enrichment analyses identified HIF-1α as an important transcriptional regulator controlling glycolytic reprogramming and terminal T cell exhaustion in response to mitochondrial insufficiency. This molecular feedback loop between mitochondrial dysfunction, ROS production and HIF-1α-mediated metabolic reprogramming represents a critical molecular checkpoint during T cell exhaustion, thus providing important translational implications for cancer immunotherapy.

Mitochondria play a pivotal role in cellular metabolism by generating energy in form of ATP. The respiratory chain complexes I-IV generate a proton gradient following OXPHOS and FAO at the inner mitochondrial membrane (IMM) that is utilized by the ATP synthase (complex V) to generate ATP from ADP and inorganic phosphate ^15, 16, 48^. In addition to their bioenergetic function, mitochondria provide intermediary metabolites controlling the fate and function of T cells by regulating signaling pathways, cellular redox balance, apoptosis and metabolic-epigenetic rewiring through post-translational DNA and histone modifications ^33, 48, 54^. Recent studies revealed that mitochondrial insufficiency, characterized by membrane depolarization, metabolic alterations and the accumulation of ROS, is tightly correlated with the functional exhaustion of T cells ^23, 24, 25, 26, 28^. Yet, a key challenge in cellular metabolism is to delineate if these changes are the cause or a consequence of T cell exhaustion. By combining novel genetic models of mitochondrial insufficiency with single-cell and functional analyses, we here show that metabolic remodelling is sufficient to elicit the exhaustion-associated gene expression program and dysfunction in Tpex cells. These data further demonstrate that mitochondrial deterioration is a cell-autonomous driver of T cell dysfunction and not merely its consequence. Mitochondrial insufficiency in mPiC-deficient T cells caused their functional exhaustion even in the absence of continued antigen exposure, suggesting that a decline in mitochondrial respiration acts – in analogy to Knudson’s “two-hit” theory in cancer ^55^ – as a second hit to initiate the exhaustion-associated gene expression program in T cells.

To establish direct causality between mitochondrial dysfunction and T cell exhaustion, we ablated the *mitochondrial phosphate carrier* (mPiC), encoded by the *Slc25a3* gene ^34, 35^. Because mPiC-dependent phosphate uptake is rate-limiting for mitochondrial ATP production ^34, 35^, *Slc25a3*^fl/fl^*Cd4*^Cre^ mice are an excellent genetic tool to investigate the cell autonomous metabolic and molecular determinants controlling T cell exhaustion. Although several studies have linked mitochondrial insufficiency to T cell exhaustion *in vitro* ^23, 24^, during chronic viral infection ^25, 26^ and in cancer ^27, 28, 29^, we here provide first-time genetic evidence that impaired mitochondrial respiration is not merely a consequence of exhaustion but, instead, promotes the transcriptional, phenotypic and functional hallmarks of T cell dysfunction. Supporting the notion that mitochondrial decline acts as a “second hit” for terminal T cell differentiation, mPiC-deficient Tpex cells did not only show an accelerated differentiation into terminally exhausted T cells during chronic viral infection, but also showed an exhaustion-associated gene expression signature and dysfunction under circumstances that are not expected to cause exhaustion, such as in acute viral infection or T cell differentiation *in vitro*.

A simple explanation for the exhaustion in response to mitochondrial deterioration in mPiC-deficient T cells could be a paucity in ATP, which was, however, not the case. mPiC-deficient T cells compensated their mitochondrial deficit by a ‘Warburg effect-like’ metabolic switch to aerobic glycolysis, thus preventing a fatal bioenergetic crisis. Glycolytic reprogramming may also represent an anabolic adaption mechanism of exhausted T cells to fuel the biosynthesis of macromolecules and to support clonal expansion and effector function ^16, 48^. However, when glucose is primarily converted into lactate to compensate for the defective mitochondrial ATP production, intermediary metabolites of the glycolytic pathway become limiting in biochemical reactions that generate reduction equivalents, such as NADPH in the oxidative branch of the *pentose phosphate pathway* (PPP) ^16, 38^. Given the crucial role of NADPH as a cellular antioxidant ^38, 48^, it is conceivable that exhausted T cells experience substantial oxidative stress. In line with recent studies ^23, 24, 28^, we found that virally and genetically exhausted T cells accumulate pathogenic levels of mitochondrial and cytosolic ROS. Scavenging excessive ROS production in mPiC-deficient T cells attenuated the expression of exhaustion markers and restored their effector function, demonstrating that ROS is not solely a by-product of defective mitochondrial respiration but a molecular rheostat linking metabolic performance to T cell exhaustion.

A previous report by Scharping *et al.* demonstrated that hypoxia and continuous antigenic stimulation together promote T cell exhaustion through Blimp-mediated repression of PCG-1α and mitochondrial dysfunction ^23^. In this study, hypoxia-induced mitochondrial ROS and chronic antigenic stimulation elicited the exhaustion-associated gene expression via the NFAT signaling pathway ^23^. In our model, T cell exhaustion occurred even in the presence of oxygen, suggesting that defective mitochondrial respiration acts as a cell-autonomous trigger for T cell exhaustion, independent of hypoxic exposure and other exogenous metabolic stressors.

Although NFAT promotes T cell exhaustion and its activity is fine-tuned by cellular ROS signaling ^18, 41, 42, 45, 46^, ablation of NFATc1 did not halt terminal differentiation of mPiC-deficient T cells. Using pathway and transcription factor enrichment analyses, we identified HIF-1α as an alternative, redox-regulated driver of T cell exhaustion. PHD-mediated hydroxylation targets HIF-1α protein for proteasomal degradation in the presence of oxygen. However, oxidation of PHD’s catalytic center by excessive cellular ROS levels leads to a stabilization of HIF-1α protein, even in presence of ambient oxygen ^47, 48^. Supporting our hypothesis that HIF-1α facilitates terminal T cell exhaustion in response to excessive ROS production, ablation of HIF-1α alleviated terminal differentiation of both genetically and virally exhausted T cells. Our findings are consistent with previous studies that revealed reduced expression of exhaustion-associated proteins in HIF-1α-deficient T cells ^56, 57, 58^, whereas stabilization of HIF-1α through genetic inactivation of its negative regulators (i.e., PHD and Von-Hippel Landau tumor suppressor) increased the expression of co-inhibitory receptor ^59, 60, 61, 62^. These data suggest that HIF-1α contributes – at least in part – to T cell dysfunction during chronic antigenic stimulation by promoting the transition of Tpex into terminally exhausted T cells.

The most prominent HIF-1α gene expression signature across different cell types and tissues involves glucose metabolism ^63, 64, 65^. Likewise, single-cell transcriptomics of virally exhausted T cells also associated the HIF-1α-dependent gene signature with glucose metabolism, including several rate-limiting enzymes of the glycolytic pathway. In addition, HIF-1α signaling may also control the expression of co-inhibitory receptors directly, including the Tim-3 ^58^. Given the elevated expression of HIF-1α and its target genes in Tpex and transitory T cells during chronic viral infection, we speculate that HIF-1a signaling initiates terminal T cell differentiation via transcriptional-metabolic reprogramming from mitochondrial respiration to aerobic glycolysis. This notion is supported by recent studies demonstrating a decisive role of TGF-β in maintaining Tpex cells ^25, 66, 67^. TGF-β inhibits the transition of Tpex into terminally differentiated Tex cells by suppressing *mechanistic target of rapamycin* (mTOR) and glycolytic reprogramming ^25, 66, 68^. Because mTOR also upregulates HIF-1a expression in activated T cells ^57, 64, 66^, we hypothesize that TGF-β maintains the stemness of Tpex cells by restricting their glucose metabolism via the AKT–mTOR–HIF-1a signaling axis. Additional evidence that moderating the glycolytic capacity of exhausted T cells sustains their stemness comes from studies using pharmacological inhibitors ^69, 70, 71, 72, 25, 73^. We show that short-term treatment with the glucose mimetic *2-desoxyglucose* reduced the expression of terminal differentiation markers and increased the persistence of CAR T cells in models of chronic viral infection and cancer immunotherapy. These findings are consistent with recent reports demonstrating that limiting glycolysis directly ^69, 70, 71^ or indirectly by inhibiting AKT ^72^, mTOR ^25^ or calcium signalling ^73^ increased the longevity and effector function of T cells.

Collectively, our findings provide a rationale for the manufacturing of exhaustion-resistant T cell products for clinical use. Because mitochondrial respiration is pivotal for the stemness of exhausted T cells, pharmacological and genetic approaches to optimize their mitochondrial endurance and/or restricting their glycolytic metabolism are promising metabolic intervention strategies to maintain (or reinvigorate) their functionality during cancer immunotherapy. Furthermore, costimulatory CAR domains favoring mitochondrial respiration over glycolysis also increased the longevity of engineered T cells, demonstrating that metabolic fine-tuning to augment (CAR) T cell stemness is a promising strategy to improve cancer immunotherapy.

## ACKNOWLEDGEMENTS

We thank Jeffery D. Molkentin (Cincinnati Children’s Hospital Medical Center, OH, USA), Anjana Rao (La Jolla Institute for Immunology, CA, USA) and Tobias Bald (University Hospital Bonn, Germany) for providing *Slc25a3*^fl/fl^ mice, CAR constructs and MC38 cells, respectively. This work was funded by the Deutsche Forschungsgemeinschaft (DFG, German Research Foundation) SFB-TR 124/3 (“FungiNet”), project number 210879364, SFB 1526/1 (“PANTAU”), project number 454193335, SFB 1583/1 (“DECIDE”), project number 492620490 and individual grants VA882/2-1 and VA882/3-2 (to M.V.). Further support was provided by the SFB 1525/1 (“Cardio-Immune-Interfaces”), project number 453989101 (to M.V., A.Z.-M., W.K., G.G. and J.D.) and SFB-TR 338/1 (“LETSimmun”), project number 452881907 (to. M.V. G.G., W.K., S.T. and S.K.). Further support came from the Interdisciplinary Center for Clinical Research (supporting the Core Units FACS and SysMed) and by the Bavarian ministry of economic affairs, regional development and energy within the project „single cell analysis in personalized medicine“ at the Helmholtz-Institute for RNA-based Infection Research (HIRI). We would like to thank Camila Takeno Cologna, Wesley Vermaelen, Anton Willems (VIB Metabolomics Expertise Center), Thorsten Bischler (IZKF Würzburg) and Fabian Imdahl (HIRI) for excellent technical assistance.

## AUTHOR CONTRIBUTION

Participated in research design: H.W., A.Ö., S.M.H., K.K., G.G., J.D., W.K., S.K. and M.V. Conducted experiments: H.W., X.Z., G.F.G., W.S., S.M.H., M.E., A.M.M., A.Ö. Performed data analysis: M.V., R.D-L., H.W., S.M.H., B.G. Provided critical research materials: S.T., G.G., A. Z-M., S.K. Wrote the manuscript: M.V. and H.W.

## METHODS

### Mice

All mice were bred and maintained under specific pathogen free conditions in the Center for Experimental Medicine (ZEMM) or the Institute for Systems Immunology at the Julius-Maximilians University of Würzburg. Mice were maintained on a 12/12 h light/dark cycle at between 20-24 °C in individually ventilated cages. Mice had access to standard chow (Ssniff; cat# V1534) and autoclaved water *ad libitum* and health status of the animals was inspected daily by the responsible caretakers. Hygiene status of the sentinel mice was monitored quarterly according to the FELASA guidelines. Both male and female mice between 8 and 24 weeks of age at the time of the experiment were used for the *in vitro* experiments described in this study. For chronic LCMV infection models, male mice between 8 and 12 weeks of age were used. All animal protocols were approved by government of Lower Franconia, Germany. C57BL/6 (JAX strain 000664), CD45.1^+^ (strain 002014), P14 (strain 037394), OT-1 (strain 003831), Ubi-GFP (strain 004353), tdTomato ^74^, mito-Dendra2 (strain 018397), *Hif1a*^fl/fl^ (strain 007561), *Tfam*^fl/fl^ (strain 026123), *Rag1*^−/–^ (strain 002216) and *Cd4*^Cre^ mice (strain 017336) were purchased from the Jackson Laboratories (JAX) and/or maintained at our institution. *Slc25a3*^fl/fl^ (mPiC) mice ^34^ and *Nfatc1*^fl/fl^ mice ^75^ were kindly provided by Jeffery D. Molkentin (Cincinnati Children’s Hospital Medical Center, OH, USA) and Anjana Rao (La Jolla Institute for Immunology, CA, USA). All animals used in this study were on a C57BL/6 genetic background.

### In vitro T cell cultures and cell lines

For *in vitro* cultures, murine CD8^+^ T cells from male and female mice (see above) were isolated from single cell suspension of lymph nodes and spleen by negative selection using the MojoSort Mouse CD8 T cell isolation kit (BioLegend). Alternatively, CD8^+^ T cells were labelled with anti-CD8α-APC (Biolegend, clone 53-6.7) and positively enriched using anti-APC microbeads (Miltenyi Biotech). T cells were cultured in modified RPMI 1640 medium with physiological glucose concentration (100 mg/dL) by diluting standard RPMI 1640 medium (Gibco) with glucose-free RPMI medium (Roth). The medium was supplemented with 10% FBS (Sigma), 50 μM 2-mercaptoethanol (β-ME), 1% penicillin/streptomycin and 1% GlutaMAX-I (all Gibco), unless otherwise stated. Platinum-E retroviral packaging cell line (Cell Biolabs Inc.) was cultured in standard DMEM with 10% FBS (Sigma) and 1% penicillin and streptomycin (Gibco) at 37°C with 5% CO_2_. MC38^OVA^ and MC38^hCD^^19^ colon carcinoma cells were grown in DMEM with 10% FBS (Sigma), 2 mM glutamine, 1 mM sodium pyruvate, 1% penicillin/streptomycin, 2 mM Hepes and 0.1 mM non-essential amino acids (all Gibco) at 37°C with 5% CO_2_.

### T cell activation and differentiation

To activate and differentiate murine CD8^+^ T cells into cytotoxic T cells (CTLs), delta-surface plates (Nunc) were pre-coated with 12 μg/ml polyclonal anti-hamster IgG (MP Biomedicals) for 2 h and washed once with PBS. In 12-well plates, 2×10^6^ cells were activated with 0.5 μg/ml of anti-CD3 (clone 145-2C1) together with 1 μg/ml anti-CD28 (clone 37.51, both Bio X Cell). After 2 days of activation, T cells were re-plated with fresh medium containing 10 ng/ml rhIL-2 or a combination of 10 ng/ml IL-7 and IL-15 (all Peprotech) and differentiated at a density of 5×10^5^ cells/ml for additional 4 days. For chronic antigenic stimulation *in vitro*, 2×10^6^ cells were activated with 0.5 μg/ml of anti-CD3 and 1 μg/ml anti-CD28 for 2 days followed by anti-CD3 stimulation. 5×10^5^ cells/ml were restimulated every other day using 0.5 μg/ml of plate bound anti-CD3 over 6 days. In some experiments, 10 µM N-acetylcysteine (NAC) or 2 mM 2-desoxyglucose (2-DG, both Sigma) were added during cultivation. T cell differentiation was performed in normoxia (21% oxygen) or in hypoxia (2% oxygen) as indicated in the figure legends.

### LCMV production and infection models

LCMV Armstrong (LCMV^ARM^) and clone 13 (LCMV^CL13^) viral stocks were produced with L929/BHK cells. Cells were infected with LCMV in a dose of 0.1 IU/cell for 1 h at RT, before the supernatant was aspirated and replaced. After 48 h and 72 h at 37 °C, the virus-containing supernatant was harvested. For the titration of viral stocks and the measurement of viral loads the organs, a limiting dilution assay was performed. 4×10^6^ cells/ml MC57G cells were seeded in 96-well plates and 50 ml of serial virus dilutions from 2×10^-2^ to 2×10^-6^ were added in eight replicates. Medium was exchanged after 48 h of incubation and after 72 h cells were washed and permeabilized/fixed using Cytofix/Cytoperm kit (BD Biosciences). Cells were stained with VL-4 hybridoma cell line supernatant containing LCMV NP-specific antibodies for 60 min at RT. As secondary antibody an anti-rat IgG AF488 (Invitrogen) was used. Virus titers were calculated by counting wells with infected cells using Axio Vert.A1 epifluorescence microscope (Zeiss). Prior to LCMV infection (d-1), mice were injected intraperitoneally (i.p.) with 200 µg anti-CD4 monoclonal depletion antibody (clone GK1.5, Bio X Cell). For LCMV^ARM^ infections, mice were infected i.p. with an infectious dose of 2×10^5^ IU. LCMV^CL13^ virus stocks were diluted in sterile PBS before intravenous (i.v.) injection of an infectious dose of 4×10^6^ IU.

### Adoptive T cell transfers

Purified P14 CD8^+^ T cells were transferred i.v. in sterile PBS into recipient mice. Depending on the experiment, a total 3-6×10^3^ P14 T cells were transferred 1 d before infection. In competitive co-transfer experiments, GFP and tdTomato-expressing WT and transgenic P14 T cells were mixed in a 1:1 ratio before transfer. In some cases, GFP^+^ andTomato^+^ P14 T cells were also labeled with cell trace violet (Thermo Fischer Scientific) before transfer. 2-DG-treated and control P14 T cells were activated with 0.5 μg/ml of anti-CD3 (clone 145-2C1) and 1 μg/ml anti-CD28 (clone 37.51, both Bio X Cell) in presence of 10 ng/ml rhIL-2 (Peprotech) and cultivated for 3-4 days *in vitro*. 1×10^6^ control and 1×10^6^ 2-DG-treated P14 T cell were co-transferred i.v. into LCMV^CL13^ infected mice 7 days post infection (d.p.i.).

### Tumor models

Ovalbumin (OVA) and human CD19-expressing MC38 colon adenocarcinoma (MC38^OVA^ and MC38^hCD^^19^) cells were grown at 37°C and 5% CO2 in standard DMEM medium supplemented with 10% FBS (Sigma), 2 mM glutamine, 1 mM sodium pyruvate, 1% penicillin/streptomycin, 2 mM Hepes and 0.1 mM non-essential amino acids (NEAA, all Gibco). 5×10^5^ MC38^OVA^ cells were subcutaneously (s.c.) injected into the flanks of 8-12 weeks old C57BL/6 mice. Six days after tumor inoculation, mice were irradiated sublethally (6 Gy). The following day, mice were injected intravenously (i.v.) with 2×10^6^ OT-I CD8^+^ T cells that were isolated from the spleen and peripheral lymph nodes of OT-1 mice and stimulated with 0.5 μg/ml anti-CD3 (clone 145-2C1), 1 μg/ml anti-CD28 antibodies (clone 37.51, both Bio X Cell) and 10 ng/ml rhIL-2 (Peprotech) for 72 h with or without 1 mM 2-desoxyglucose (2-DG, Sigma). Tumor size was assessed every day using a caliper. For some experiments, tumor tissue was harvested digested with 1 mg/ml collagenase type I and 100 μg/ml DNase I (both Roche) and T cell infiltration analyzed by flow cytometry. Alternatively, *Rag1*^−/–^ mice were s.c. injected with 1×10^6^ MC38^hCD19^. After 7 days, 2×10^6^ anti-hCD19 CAR T cells ^42^ treated with and without 2 mM 2-DG for 2 days were adoptively transferred i.v.

### Flow cytometry

Flow cytometric staining was performed as previously described (Vaeth et al., 2019). Briefly, cells were stained with Fixable Viability Dye eFluor 780 (eBioscience) for 10 min in PBS at RT together with an anti-FcgRII/FcgRIII antibody (clone 2.4G2; Bio X Cell) to prevent unspecific binding. After washing, surface antigens were stained with fluorophore-conjugated antibodies in PBS containing 0.5% BSA for 20 min at RT in the dark. For intracellular cytokine staining, cells were stimulated with 1 μM ionomycin (BioMol) and 30 nM phorbol-12-myristat-13-acetate (PMA, Sigma) or with 1 μg/ml of anti-CD3 (clone 145-2C1) together with 1 μg/ml anti-CD28 (clone 37.51, both Bio X Cell) in the presence of 2 μg/ml brefeldin A and 2 μM monensin (both eBioscience) for 4 h at 37°C. After surface staining, cells were fixed with IC-fixation buffer (eBioscience) and intracellular cytokines were stained using 1x permeabilization buffer (both eBioscience) for 40 min at RT. A complete list of flow cytometry antibodies can be found in Supplementary Table S2. For the quantification of the mitochondrial content and membrane potential, T cells were loaded with 500 nM MitoTracker DeepRed and 2 nM Tetramethylrhodamin-Ethylester (TMRE) (both Invitrogen). As control for TMRE, cells were pre-treated with 20 µM Trifluoromethoxy carbonylcyanide phenylhydrazone (FCCP, Cayman Chemicals) for 15 min to depolarize the membrane potential. For intracellular ROS determination, MitoSOX and CellROX fluorescent dyes (both ThermoFischer Scientific) were diluted in HBSS (Gibco) buffer and culture medium, respectively. Cells were incubated with the probes for 30 min at 37°C and washed twice with PBS before analysis. To analyze T cell proliferation, T cells were loaded with 5 μM CellTrace Violet (ThermoFischer Scientific) according to manufacturer’s instructions. To measure HIF-1α protein expression, stimulated T cells were fixed with the Foxp3/TF staining buffer set (eBioscience) for 30 min at 4°C and stained intranuclearly with rabbit polyclonal anti-mouse HIF-1α (Cell Signaling) o/n in permeabilization buffer (eBioscience). After washing, the primary antibody was detected using a donkey polyclonal anti-rabbit IgG secondary antibody conjugated to Alexa Fluor 647 (Invitrogen) diluted 1:1000 in permeabilization buffer for 30 min. All sample acquisition was performed with BD Celesta (BD Biosciences) or Attune Nxt (ThermoFischer Scientific) flow cytometers and further analyzed with the FlowJo software (Tree Star).

### Retroviral transduction

Retroviral infection of primary T cells was performed as described previously (Vaeth et al., 2017a). Briefly, Platinum-E retroviral packaging cells were transfected transiently with modified pMIG retroviral plasmids (Addgene, #9044) or a second-generation anti-hCD19 CAR construct (MSCV-myc-CAR2A-Thy1.1) ^42^ using the GeneJet reagent (SignaGene). The transfection medium was replaced 24 h later with standard DMEM medium and the supernatant containing retroviral particles was collected 2 and 3 days after transfection. CD8^+^ T cells were isolated by negative selection using the MojoSort Mouse CD8^+^ T cell isolation kit (BioLegend) and cultivated as described above. 24 h after activation, the medium of the T cells was replaced by retroviral supernatant and the cells were transduced by spin-infection (2.500 rpm, 30°C, 90 min) in the presence of 10 mg/ml polybrene (SantaCruz). After transduction, cells were incubated at 37°C for 4 h before the viral supernatant was removed and replaced by fresh medium. After 2 days of stimulation, transduced T cells were rested for additional 2 days in fresh medium containing 10 ng/ml rhIL-2 (Peprotech) before 1×10^5^ pMIG-empty vector (Ametrine^+^) and 1×10^5^ pMIG-SLC25A3 (GFP^+^) expressing P14 T cells were co-transferred into LCMV^CL13^ infected recipient mice or as described above. T cells transduced with the anti-hCD19 CAR construct ^42^ were cultivated for additional 2 days in presence or without 2 mM 2-DG (Sigma) before adoptive transfer into MC38-hCD19 tumor-bearing mice.

### Mitochondrial turnover measurements

Mito-Dendra2 mice ^32^ were infected with LCMV^CL13^ as described above. 14-16 days post infection, splenocytes were harvested and lilluminated with LED blue light using a long-pass filter (∼ 405 nm) for 30 seconds to photoconvert the mitochondrially localized Dendra2 protein. Afterwards, splenocytes were labelled with cell trace violet (CTV) (ThermoFischer Scientific) and cultured for 48 h with 10 ng/ml rhIL-2 (Peprotech) *in vitro*. Mitochondrial regeneration capacity was calculated based on the reduction of geometric mean fluorescence intensity (gMFI) value of mito-Dendra2-red: (gMFI_0h_-gMFI_48h_) / gMFI_0h_.

### Seahorse extracellular flux analysis

Mitochondrial respiration and lactate secretion of T cells was measured as their oxygen consumption rate (OCR) and glycolytic proton efflux rate (glycoPER), respectively, using an oxygen-controlled XFe96 extracellular flux analyzer (Seahorse Bioscience). XFe96 cell culture microplates (Agilent) were pre-coated with 22 μg/ml Cell-Tak (Corning) and 1-2×10^5^ T cells per well were attached in 5-8 replicates in Seahorse XF RPMI medium (Agilent) supplemented with 2 mM L-glutamine (Gibco), 1 mM sodium pyruvate (Sigma) and 10 mM D-glucose (Sigma). After 1h incubation in a CO_2_-free incubator at 37°C, glycolytic and mitochondrial stress tests were performed according to the manufacture’s recommendation. In brief, for assessing glycolysis, basal extracellular acidification rate (ECAR) was measured followed by addition of 0.5 μM rotenone (AdipoGen) and 0.5 μM antimycin A (Sigma) to inhibit mitochondrial complex 1 and 3, respectively. At the end of the measurement, 50 mM 2-DG (Sigma) was added to completely block glycolysis. To analyze mitochondrial respiration, basal oxygen consumption was measured followed by the addition of 2 μM oligomycin (Cayman Chemicals), an ATP synthase inhibitor, 1 μM of the protonophore carbonyl cyanide-*4*-(trifluoromethoxy)-phenylhydrazone (FCCP, Cayman Chemical) and 0.5 μM rotenone (AdipoGen) together with 0.5 μM antimycin A (Sigma). The basal oxygen consumption was calculated by subtracting the OCR after rotenone and antimycin A treatment from the OCR before oligomycin treatment. The maximal OCR was calculated by subtracting the OCR after rotenone and antimycin A treatment from the OCR measured after addition of FCCP.

### Metabolomic profiling

To analyze ^12^C and ^13^C polar metabolites, CD8^+^ T cells were activated in RPMI containing 1 g/l ^12^C-glucose as described above. After 2 days of stimulation and resting o/n in fresh medium containing 10 ng/ml rhIL-2 (Peprotech), T cell were labelled in modified RPMI 1640 containing 1 g/l ^13^C_6_-glucose (Sigma). After washing with pre-warmed PBS, metabolite extraction was performed in ice-cold 80% methanol containing 1 μM of ^13^C5-d5-^15^N glutamic acid, 1 μM d7-^15^N4 arginine, 1 μM d27 myristic acid and 1 μM d12 glucose as internal standards. Following centrifugation with 20.000 g for 10 min at 4°C, the supernatant containing polar metabolites was transferred to a new tube and stored at −80°C. The pellet was used for protein quantification as an internal normalization method. 10 μl sample lysate was loaded into a Dionex UltiMate 3000 LC System (Thermo Scientific) equipped with a C-18 chromatography column (Acquity UPLC-HSS T3 1. 8 μm; 2.1 x 150 mm, Waters) coupled to a Q Exactive Orbitrap mass spectrometer (Thermo Scientific) operating in negative ion mode. A step gradient was carried out using solvent A (10 mM TBA and 15 mM acetic acid) and solvent B (100% methanol). The gradient started with 0% of solvent B and 100% solvent A and remained at 0% B until 2 min post injection. A linear gradient to 37% B was carried out until 7 min and increased to 41% until 14 min. Between 14 and 26 minutes the gradient increased to 100% of B and remained at 100% B for 4 minutes. At 30 min the gradient returned to 0% B. The chromatography stopped at 40 min. The flow was kept constant at 250 μl/min and the column was placed at 25°C throughout the analysis. The MS operated in full scan mode (m/z range: [70–1050]) using a spray voltage of 3.2 kV, capillary temperature of 320 °C, sheath gas at 40.0, auxiliary gas at 10.0. The AGC target was set at 3e6 using a resolution of 140.000, with a maximum IT fill time of 512 ms. Data collection was performed using the Xcalibur software (Thermo Scientific) and analyzed by integrating the peak areas using the El-Maven to Polly data processing engine (Elucidata). Cellular sphingolipids were analyzed after extraction with methanol:chloroform (2:1) using a 1290 Infinity II HPLC coupled with a 6495C triple-quadrupole mass spectrometer (Agilent Technologies) as previously described (Naser et al., 2020). Alternatively, water-soluble metabolites were extracted with 500 μL ice-cold MeOH/H_2_O (80/20, v/v) containing 0.01 μM lamivudine and 10 µM each of D4-succinate, D5-glycine, D2-glucose and 15N-glutamate as standards (Sigma-Aldrich). After centrifugation of the resulting homogenates, supernatants were transferred to an RP18 SPE (50 mg/1 mL tubes, Phenomenex) that had been activated with 0.5 mL CH_3_CN and conditioned with 0.5 mL of MeOH/H2O (80/20, v/v). The eluate of RP18 SPE-column was evaporated in a SpeedVac (Thermo Fisher Scientific). Dry sample extracts were re-dissolved in 100 μL 5 mM NH_4_OAc in CH_3_CN/H_2_O (50/50, v/v). 15 μL supernatant was transferred to LC-vials. For LC/MS analysis 3 μL of each sample was applied to a XBridge Premier BEH Amide (2.5 μm particles, 100 x 2.1 mm) UPLC-column (Waters). Metabolites were separated with Solvent A, consisting of 5 mM NH_4_OAc in CH_3_CN/H_2_O (40/60, v/v) and solvent B consisting of 5 mM NH_4_OAc in CH3CN/H2O (95/5, v/v) at a flow rate of 200 μL/min at 45°C by LC using a DIONEX Ultimate 3000 UHPLC system (Thermo Fisher Scientific). A linear gradient starting after 2 min with 100% solvent B decreasing to 0% solvent B within 23 min, followed by 17 min 0% solvent B and a linear increase to 100% solvent B in 1 min. Recalibration of the column was achieved by 7 min pre-run with 100% solvent B before each injection. HPLC-MS solvents, NH_4_OAc, standards and reference compounds were purchased from Merck. MS-analyses were performed on a high-resolution Q Exactive mass spectrometer equipped with a HESI probe (Thermo Fisher Scientific) in alternating positive and negative full MS mode with a scan range of 69.0–1000 m/z at 70K resolution and the following ESI source parameters: sheath gas: 30, auxiliary gas: 1, sweep gas: 0, aux gas heater temperature: 120°C, spray voltage: 3 kV, capillary temperature: 320°C, S-lens RF level: 50. XIC generation and signal quantitation was performed using TraceFinder V 3.3 (Thermo Fisher Scientific) integrating peaks which corresponded to the calculated monoisotopic metabolite masses (MIM +/− H+ ± 3 mMU).

### Bulk RNA-sequencing

For bulk RNA-sequencing (RNA-seq), naive T cells of *Slc25a3*^fl/fl^*Cd4*^Cre^ or WT mice were left unstimulated, stimulated with anti-CD3/CD28 for 2 days or differentiated for 6 days into CTLs cells. For each condition, 1×10^6^ cells were directly harvested in RNAprotect Cell reagent and stored at −80°C o/n before total RNA was extracted using the RNeasy Plus Micro Kit (both Quiagen). RNA quality and quantity were checked using a 2100 Bioanalyzer with the RNA 6000 Nano kit (Agilent Technologies) and only samples with RIN > 7 were used. cDNA libraries suitable for sequencing were prepared from 500 ng of total RNA with TruSeq mRNA Stranded Library Prep Kit from Illumina according to manufacturer’s instructions (1/2 volume). Libraries were quantified by QubitTM 3.0 Fluometer (Thermo Scientific) and quality was checked using 2100 Bioanalyzer (Agilent) with High Sensitivity DNA kit (Agilent). 0.5 ng of each library was subjected to a tagmentation-based protocol (Nextera XT, Illumina) using a quarter of the recommended reagent volumes. Libraries were quantified again by QubitTM 3.0 Fluometer (Thermo Scientific) and quality was checked using 2100 Bioanalyzer with High Sensitivity DNA kit (Agilent) before pooling. In both experiments, sequencing of pooled libraries, spiked with 1% PhiX control library, was performed at 19-36 million reads/sample in single-end mode with 75 nt read length on the NextSeq 500 platform (Illumina) with 1 High Output Kit v2.5. Demultiplexed FASTQ files were generated with bcl2fastq2 v2.20.0.422 (Illumina). To assure high sequence quality, Illumina reads were quality and adapter trimmed via Cutadapt [1] version 2.5 using a cutoff Phred score of 20 in NextSeq mode and reads without any remaining bases were discarded. Processed reads were subsequently mapped to the mouse genome (GRCm38.p6 primary assembly and mitochondrion) using STAR v2.7.2b with default parameters based on RefSeq annotation version 108.20200622 for GRCm38.p6 [2]. Read counts on exon level summarized for each gene were generated using featureCounts v1.6.4 from the Subread package [3]. Multi-mapping and multi-overlapping reads were counted non-strand-specific with a fractional count for each alignment and overlapping feature. The count output was utilized to identify differentially expressed genes using DESeq2 [4] version 1.24.0. Read counts were normalized by DESeq2 and fold-change shrinkage was applied by setting the parameter “betaPrior=TRUE”. Differences in gene expression were considered significant if padj < 0.05. The DEseq2 data was further analyzed by gene set enrichment analysis (GSEA) and visualized in Cytoscape using the plugin ErichmentMap in edge cutoff 0.5. Pathways were filtered and displayed when p value < 0.005 and Q value < 0.1. A complete list of all gene sets used in this study can be found in Supplementary Table S1.

### Single-cell RNA sequencing

CD8^+^ PD-1^+^ CD44^+^ (Figure 1) or CD8^+^ tdTomato^+^ (HIF-1α-deficient) and CD8^+^ GFP^+^ tdTomato^+^ (WT) P14 T cells (Figure 5) were FACS-sorted from the spleen or LNs of chronically infected mice using a FACSAria III cell sorter (BD Biosciences). To multiplex different samples for scRNA sequencing, single-cell suspensions after FACS sorting were labelled with different TotalSeqA antibodies (Biolegend). Each sample was labelled with one specific hashtag (1:400 dilution) and CITE-seq antibodies (1:800 dilution, all Biolegend). After labelling for 30 min at 4 °C, samples were washed three times and multiplexed in equal cell numbers at a density of 10^6^ cells/ml. Singe cells were encapsulated into droplets with the Chromium^TM^ Controller (10x Genomics) and processed following manufacturer’s specifications. Transcripts captured in all the cells encapsulated with a bead were uniquely barcoded using a combination of a 16 bp 10x Barcode and a 10 bp unique molecular identifier (UMI). cDNA libraries were generated using the Chromium Single Cell 3’ Library & Gel Bead Kit v2 (10x Genomics) following the detailed protocol provided by the manufacturer. Libraries were quantified by QubitTM 3.0 Fluometer (ThermoFisher) and quality was checked using 2100 Bioanalyzer with High Sensitivity DNA kit (Agilent). Libraries were sequenced with the NovaSeq 6000 platform (S1 Cartridge, Illumina) in 50 bp paired-end mode. The sequencing data was demultiplexed using CellRanger software (version 2.0.2). The reads were aligned to mouse mm10 reference genome using STAR aligner. Aligned reads were used to quantify the expression level of mouse genes and generation of gene-barcode matrix. Subsequent data analysis was performed using Seurat R package (version 3.2) (Stuart et al., 2019).

### scRNA sequencing analysis

Quality control was performed, and viable cells were selected by excluding cells with number of genes lower than 500 and above 4000 and/or having more than 8% of mitochondrial transcripts (Figure 1). Alternatively, cells having less than 100 transcripts in total and/or more than 8% of mitochondrial transcripts were removed from the analysis (Figure 5). To demultiplex hashtags HTODemux function in Seurat package was used with standard parameters. Cross-sample doublet cells were detected based on hashtag signal (TotalSeq A0301 and A0302). Cells that were classified as ‘singlet’ and identified by hashtags were retained and used for downstream analysis. Principle component (PC) analysis was used for dimensionality reduction and a uniform manifold approximation and projection (UMAP) was performed on the first 15 PC dimensions. Cells were assigned to clusters with the FindNeighbors function in Seurat on the same PC dimensions and the UMAP clusters were identified using the function FindClusters with a resolution of 0.5 and 0.4 in Figure 1 and Figure 5, respectively. Contaminating cell types or proliferating cell were removed from the analysis based on marker genes or *Mki67* gene expression, respectively. WT and HIF-1α-deficient samples (Figure 6) were identified by TotalSeq A0301 and A0302 Hashtag-antibodies, respectively. Enrichment scores for *mitochondrial respiration chain complex assembly* (gene set M11099), *pentose phosphate pathway* (M1386) and HIF-1α target gene signature (combined M17905 and M2513) were calculated using gene lists with AddModuleScore function from Seurat. Differentially expressed genes between WT and HIF-1α-deficient P14 T cells were identified with the FindMarkers function in Seurat (min.pct = 0.25 and logfc.threshold = 0.25) and then subjected to pathway analysis using the EnrichR website (https://maayanlab.cloud/Enrichr/). The KEGG Pathway 2021 Human database was used as a reference to test for pathway enrichment with standard EnrichR settings. Adjusted p-values (padj) comparing gene expression of clusters and samples were calculated with the FindMarker function in Seurat; other p-values were calculated using Fisher’s exact test.

### Trajectory analysis and gene expression visualization

Trajectories were predicted using Slingshot 1.4.0 package, using function slingshot with a threshold of 0.001, a stretch of 2 and starting with cluster ‘‘Tpex1’’ based on *Sell* (CD62L) expression ^10^. Gene expression was visualized as heat maps, volcano and violin plots using the Seurat package and ggplot2.

### Immunoblotting

Cell lysates were prepared in RIPA buffer supplemented with complete protease-inhibitor cocktail (Roche) and mixed in equal amounts with Laemmli buffer (Roth) before denaturation at 98 °C for 5 min. Protein quantification was performed with the Pierce 660 nm Protein Assay (Pierce). Protein samples were separated via SDS-PAGE, blotted onto nitrocellulose membranes, blocked with 5% BSA and incubated with monoclonal rabbit anti-HIF-1α (Cell Signaling Technology, clone D1S7W). Detection was carried out with a goat anti-rabbit horse radish peroxidase (HRP) conjugated secondary antibody (BioRad), visualized with the enhanced chemiluminescent SuperSignal reagent (Pierce). For the detection of PGC-1α, TFAM and elector transport chain (ETC) complex expression, Tpex and Tex cells were FACS sorted from LCMV^CL13^ infected mice 30 d.p.i. and 5×10^5^ T cells were directly lysed in RIPA buffer. Detection of PGC-1α, TFAM and ETC complexes was carried out using rabbit anti-PGC-1α (Abcam), rabbit-anti-TFAM (Abcam) and an OxPhos rodent antibody cocktail (Thermo) with goat anti-mouse or rabbit HRP-conjugated secondary antibodies (both BioRad).

### Statistical Analysis

The results are shown as mean ± standard error of the means (SEM). To determine the statistical significance of the differences between the experimental groups unpaired or paired Student’s t tests, 2-way ANOVA and Mantel-Cox tests were performed using the Prism 9 software (GraphPad). Sample sizes were based on experience and experimental complexity, but no methods were used to determine normal distribution of the samples. Differences reached significance with p values < 0.05 (noted in figures as *), p < 0.01 (**) and p < 0.001 (***). The figure legends contain the number of independent experiments or mice per group that were used in the respective experiments.

## Data availability

The bulk and single cell RNA sequencing datasets have been deposited at GEO under accession numbers GSE212298 (reviewer token: mbcneaccptgdbkf), GSE214003 (ibmbwgkmphsrvof) and GSE213847 (wdepmsmihlspdij). Any additional information required to analyze the data reported in this paper is available upon reasonable request. Source data are provided with this paper.

## Code availability

No original codes have been generated in this study.

## EXTENDED DATA

**Extended Data Figure 1. Transcriptional and metabolic features of T cell exhaustion. (a)** Naïve C57BL/6 mice were chronically infected with the LCMV strain clone 13 (LCMV^CL13^) and CD8^+^CD44^+^PD-1^hi^ T cells were FACS sorted and subjected to single cell (sc) RNA sequencing at day 21 post infection. **(b)** Heat map of marker gene expression associated with the uniform manifold approximation and projection (UMAP) clusters depicted in Fig. 1a. **(c)** Normalised gene expression of *Id3*, *Il7r, Cxcr5, Ccr7, Zbp1, Irf7, Id2, Lag3, Cd244a* (2B4) and *Tnfrsf9* (41-BB) projected onto UMAP clusters. **(d)** Network clustering of publicly available bulk RNA sequencing data of Tpex and Tex cells (Utzschneider *et al*. 2020; GSE142686) using significantly (p < 0.05) enriched gene expression signatures. Upregulated gene sets in Tpex and Tex are shown in orange and blue, respectively. **(e)** Gene set enrichment analysis (GSEA) of *Mootha mitochondria* (gene set M9577), *mitochondrial translation* (M27446) and *respiratory electron transport ATP synthesis by chemiosmotic coupling by uncoupling proteins* gene signatures (M1025) in Tpex and Tex cells.

**Extended Data Figure 2. Characterization of mitochondria in Tpex and Tex cells. (a)** Analysis of mitochondrial mass/volume in Tpex and Tex cells using LCMV^CL13^ infected mito-Dendra2 mice (without photoconversion). **(b)** Mitochondrial regeneration capacity of non-divided Tpex and Tex cells *ex vivo*. T cells were isolated from LCMV^CL13^ infected mito-Dendra2 mice and photo-converted with 405 nm laser light. Mito-Dendra2-red and mito-Dendra2-green was measured by flow cytometry 48 h after photoconversion. **(c)** Flow cytometric analysis of mitochondrial volume/mass and membrane potential in Tnaive, Tpex and Tex cells using MitoTracker and TMRE probes, respectively; means ± SEM of 9-11 mice. **(d)** Western Blot analyses of mitochondrial electron transport chain (ETC) complexes, PGC-1α and TFAM expression in Tnaive, Tact, Tpex and Tex cells. Respective T cell subsets were isolated from WT mice by FACS sorting 21 days after chronic LCMV infection. **(e)** Analysis of mitochondrial (ETC) complexes by flow cytometry 21 days after LCMV^CL13^ infection; means ± SEM of 4 mice. *, p<0.05; **, p<0.01; ***, p<0.001 by unpaired Student’s t-test in (c-e).

**Extended Data Figure 3. Immunological characterization of *Slc25a3*^fl/fl^*Cd4*^Cre^ mice and mPiC-deficient CD8^+^ T cells. (a)** Transport of inorganic phosphate (Pi) into the mitochondrial matrix is mediated by the mitochondrial phosphate carrier (mPiC), encoded by the *Slc25a3* gene. **(b-g)** Flow cytometric characterization of mPiC-deficient (*Slc25a3*^fl/fl^*Cd4*^Cre^) mice. **(b)** Total cell numbers of thymus, spleen and lymph nodes (LNs) of 8-14 weeks old WT and *Slc25a3*^fl/fl^*Cd4*^Cre^ mice; means ± SEM of 7-8 mice. **(c)** Representative flow cytometric analyses of thymic T cell subsets in WT and *Slc25a3*^fl/fl^*Cd4*^Cre^ mice. **(d, e)** Analysis of peripheral CD4^+^ and CD8^+^ T cell subsets (d) and Foxp3^+^ Treg cells (e) in spleens and LNs of WT and *Slc25a3*^fl/fl^*Cd4*^Cre^ mice; means ± SEM of 7-8 mice. **(f, g)** Flow cytometric analysis of CD44^−^ CD62L^+^ (naïve), CD44^+^CD62L^+^ (central memory) and CD44^+^CD62L^−^ (effector) CD4^+^ and CD8^+^T cells of WT and *Slc2a3*^fl/fl^*Cd4*^Cre^ mice; means ± SEM of 7-8 mice. **(h)** Analyses of oxygen consumption rate (OCR) of WT and mPiC-deficient (*Slc25a3*^fl/fl^*Cd4*^Cre^) cytotoxic lymphocytes (CTLs) at day 6 of culture using a Seahorse extracellular flux analyzer; means ± SEM of 3 mice. **(i)** Ratio of extracellular acidification rate (ECAR) to OCR in WT and mPiC-deficient T cells at day 3 and day 6 of culture; means ± SEM of 3 mice. **(j)** Isotope tracing of glucose-derived metabolites in WT and GLUT3-deficient T cells by liquid chromatography and mass spectrometry (LC/MS) after incubation with ^13^C-glucose. Analysis of (m+3) glycolytic and (m+2) TCA cycle metabolites in WT and mPiC-deficient T cells after 5 min and 6 h labeling with ^13^C-glucose. Fractional enrichments of ^13^C-metabolites are shown as means ± SEM of 3 biological replicates. **(k)** Analysis of ^12^C and ^13^C-labelled lactate (m+3) and citrate (m+2) by LC/MS after incubation with ^13^C-glucose for 6 h. Ratio of ^13^C-lactate to ^13^C-citrate in WT and mPiC-deficient T cells; means ± SEM of 3 biological replicates. **(l, m)** MA plots of differentially expressed genes (DEGs) in WT *versus* mPiC-deficient T cells activated for with anti-CD3/CD28 (day 2) and after differentiation into CTLs (day 6). *, p<0.05; **, p<0.01; ***, p<0.001 by unpaired Student’s t-test in (d-j).

**Extended Data Figure 4. Differentiation of WT and mPiC-deficient CD8^+^ T cells with IL-7 plus IL-15 and chronic antigenic stimulation. (a-c)** Expression of activation (a), memory (b) and exhaustion markers (c) on cytotoxic lymphocytes (CTLs) and memory-like T cells differentiated from naïve WT and mPiC-deficient (*Slc25a3*^fl/fl^*Cd4*^Cre^) T cells. CD8^+^ T cells were activated with anti-CD3/CD28 for 2 days followed by 4 days resting in IL-2 or a combination of IL-7 and IL-15 to generate CTL and memory-like T cells, respectively; means ± SEM of 3 mice. **(d-f)** Expression of activation (d), memory (e) and exhaustion markers (f) on WT and mPiC-deficient T cells differentiated under acute and chronic antigenic stimulation. CD8^+^ T cells were activated with anti-CD3/CD28 for 2 days followed by 4 days of resting in IL-2 (acute) or continuous antigenic stimulation with anti-CD3/CD28 over 6 days (chronic); means ± SEM of 6 mice. **(g)** Analysis of polyfunctional TNFα, IFNγ and IL-2 expression by WT and mPiC-deficient T cells differentiated under acute and chronic antigenic stimulation and restimulation with anti-CD3/CD28 for 6 h; means ± SEM of 3 mice. *, p<0.05; **, p<0.01; ***, p<0.001 by unpaired Student’s t-test in (a-g).

**Extended Data Figure 5.** Mitochondrial respiration controls the proliferation and clonal expansion of virus-specific T cells *in vivo*. (a) Adoptive co-transfer of naïve GFP^+^ WT and tdTomato^+^ mPiC-deficient (*Slc25a3*^fl/fl^*Cd4*^Cre^) P14 T cells into C57BL/6 mice before infection with LCMV clone 13 (LCMV^CL13^). Flow cytometric analysis of CD44, Tim-3 and Ly108 expression on donor P14 T cells in the spleen and LNs 14 days post infection (d.p.i.). **(b)** Representative flow cytometric analyses of splenic GFP^+^ (WT) and tdTomoto^+^ (mPiC-deficient) donor P14 T cells at different timepoints after transfer into LCMV^CL13^ infected mice. **(c)** Relative proportions of WT and mPiC-deficient donor P14 T cells in the spleens and LNs of chronically infected recipient mice; means ± SEM of 4-8 mice per timepoint. **(d)** Proliferation of WT and mPiC-deficient CD8^+^ T cells by CellTrace violet (CTV) dilution over a course of 4 days *in vitro*. **(e)** Cellular expansion of WT and mPiC-deficient T cells in vitro; means ± SEM of 3 mice per timepoint. **(f)** Proliferation analysis of WT and mPiC-deficient P14 T cells *in vivo*. Adoptive co-transfer CTV-labelled GFP^+^ WT and tdTomato^+^ mPiC-deficient P14 T cells into chronically infected C57BL/6 WT host mice. Flow cytometric analysis of proliferation and cellular expansion 3 days after adoptive transfer. *, p<0.05; **, p<0.01; ***, p<0.001 by paired (a and f) and unpaired (d) Student’s t-tests.

**Extended Data Figure 6. Terminal differentiation of mPiC-deficient T cells is mediated by ROS and HIF-1α. (a)** Isotope tracing of glucose-derived 6-phosphogluconate (6PG) and pentose-5-phosphate (P5P) in WT and mPiC-deficient (*Slc25a3*^fl/fl^*Cd4*^Cre^) T cells by liquid chromatography and mass spectrometry (LC/MS) after incubation with ^13^C-glucose for 24 h. Fractional enrichment of ^13^C-glucose derived metabolites in WT and mPiC-deficient T cells is depicted in grey and blue, respectively. Statistical analysis on total metabolite levels; 5 biological replicates per group. **(b)** Enrichment of pentose phosphate pathway gene expression signatures (gene set ID M1386) in UMAP clusters representing Tpex and Tex cell subsets as shown in Fig. 1a-d. **(c)** Mitochondrial ROS may enhance the Ca^2+^ / calcineurin / NFAT signaling pathway in activated T cells. **(d, e)** Differentiation of WT, mPiC-deficient (*Slc25a3*^fl/fl^*Cd4*^Cre^) and mPiC/NFATc1 double-deficient (*Slc25a3*^fl/fl^*Nfatc1*^fl/fl^*Cd4*^Cre^) T cells *in vitro*. Representative flow cytometric analysis (d) and quantification of PD-1 and Tim-3 expression on WT, mPiC-deficient and mPiC/NFATc1 double-deficient T cells (e); means ± SEM of 3 mice. **(f)** Analysis of *Hif1a* gene expression in different Tpex and Tex UMAP clusters analyzed by scRNA sequencing as shown in Fig. 1a-d. **(f)** Quantification of HIF-1α protein expression in Tpex and Tex cells in all CD8^+^ T cells and LCMV-specific (GP_33-41_ teramer^+^) T cells in the spleen of WT mice 21 days after LCMV^CL^^13^ infection; means ± SEM of 4 mice. **(h)** Enrichment analysis of the HIF-1α target gene expression (combined gene sets M17905 and M2513) in Tpex and Tex cell clusters analyzed by scRNA sequencing as shown in Fig. 1a-d. **(i)** ROS prevents HIF-1α protein degradation through oxidative inhibition of prolyl hydoxylases (PHDs). **(j)** Isotope tracing of glucose-derived TCA cycle metabolites in WT and mPiC-deficient T cells after incubation with ^13^C-glucose for 24 h. Fractional enrichment of ^13^C-glucose derived metabolites in WT and mPiC-deficient T cells is depicted in grey and blue, respectively. Statistical analysis based on total metabolite levels; 5 biological replicates per group. **(k)** Western blot analysis of HIF-1α protein expression in WT and mPiC-deficient T cells with and without ROS scavenging using N-acetyl cysteine (NAC). *, p<0.05; **, p<0.01; ***, p<0.001 by unpaired Student’s t-test in (a), (e), (g) and (j).

**Extended Data Figure 7. Ablation of TFAM in T cells causes metabolic reprogramming, ROS production, HIF-1α stabilization and terminal exhaustion. (a, b)** Analyses of oxygen consumption rate (OCR) and glycolytic proton efflux rate (glycoPER) in anti-CD3/CD28 activated WT and TFAM-deficient (*Tfam*^fl/fl^*Cd4*^Cre^) T cells using a Seahorse extracellular flux analyzer; means ± SEM of 2-3 mice. **(c)** Flow cytometric analysis of cellular ROS production (CellROX) in anti-CD3/CD28 stimulated WT and TFAM-deficient CD8^+^ T cells; means ± SEM of 4 mice. **(d)** Analysis of HIF-1α protein expression in activated WT and TFAM-deficient CD8^+^ T cells by flow cytometry; means ± SEM of 4 mice. **(e, f)** Flow cytometric detection of exhaustion (e) and stemness marker (f) expression in WT and TFAM-deficient T cells, means ± SEM of 4 mice. *, p<0.05; **, p<0.01; ***, p<0.001 by unpaired Student’s t-test in (c-f).

**Extended Data Figure 8. HIF-1α controls the exhaustion of virus-specific T cells. (a)** Adoptive co-transfer of tdTomato^+^GFP^+^ WT and tdTomato^+^ HIF-1α-deficient (*Hif1a*^fl/fl^*Cd4*^Cre^) P14 T cells into C57BL/6 mice before chronic infection with LCMV clone 13 (LCMV^CL13^). 21 days post infection (d.p.i.), donor WT and HIF-1α-deficient P14 T cells were FACS sorted, barcoded and multiplexed in a 1:1 ratio before subjected to single cell (sc) RNA sequencing. **(b)** Heat map of marker gene expression associated with the uniform manifold approximation and projection (UMAP) clusters depicted in Fig. 5a. **(c)** Normalized gene expression of *Id3*, *Il7r, Cxcr5, Id2, Lag3* and *Cd244a* (2B4) projected onto UMAP clusters. **(d)** Enrichment of mitochondrial (gene set ID M9577) and respiratory chain gene expression signatures (M19046) in UMAP clusters representing Tpex and Tex cell subsets as shown in Fig. 5a. **(e, f)** Chronic viral infection of WT and *Hif1a*^fl/fl^*Cd4*^Cre^ mice with LCMV^CL13^. **(e)** Analysis of LCMV-specific (GP_33-41_ teramer^+^) CD8^+^ T cells in the spleen and LNs of WT and *Hif1a*^fl/fl^*Cd4*^Cre^ mice 27 d.p.i.; means ± SEM of 3-5 mice. **(f)** Flow cytometric analysis of Tpex and Tex cells among CD44^+^PD-1^+^ and LCMV-specific (GP_33-41_ teramer^+^) CD8^+^ T cells in the spleen and LNs of WT and *Hif1a*^fl/fl^*Cd4*^Cre^ mice 14 d.p.i.; means ± SEM of 3-5 mice. *, p<0.05; **, p<0.01; ***, p<0.001 by unpaired Student’s t-test in (f).

**Extended Data Figure 9. HIF-1α-mediated glycolytic reprogramming drives terminal T cell differentiation. (a, b)** Differentiation of WT and HIF-1α-deficient (*Hif1a*^fl/fl^*Cd4*^Cre^) T cells under hypoxia *in vitro*. Flow cytometric analyses of exhaustion (a) and memory (b) marker expression in WT and HIF-1α-deficient T cells, means ± SEM of 4-5 mice. **(c)** Limitation of glycolysis prevents the upregulation of co-inhibitory receptors on mPiC-deficient (*Slc25a3*^fl/fl^*Cd4*^Cre^) T cells *in vitro*. Flow cytometric analyses of Tim-3, granzyme A and Lag3 expression in WT and mPiC-deficient T cells treated with or without 2-desoxy-glucose (2-DG); means ± SEM of 3 mice. **(d-g)** Inhibition of glycolytic reprogramming sustains the stemness of virus-specific T cells *in vivo*. **(d)** Adoptive co-transfer of 2-DG pre-treated (tdTomato^+^ GFP^+^) and control (GFP^+^) P14 T cells into C57BL/6 mice after chronic infection with LCMV^CL^^13^ (as shown in Fig. 6h). **(e)** Flow cytometric analysis of P14 Tpex and Tex cells 21 d.p.i.; n = 6 mice. **(f)** Ratios of 2-DG pre-treated (tdTomato^+^GFP^+^) and control (GFP^+^) donor P14 T cells in the spleens and LNs of recipient mice at days 10 and 21 after transfer; means ± SEM of 6 mice per timepoint. **(g)** Analysis of polyfunctional TNFα, IFNγ and IL-2 expression after PMA/iono re-stimulation of 2-DG-treated and control P14 T cells 21 days after co-transfer into chronically infected mice (as shown in Fig. 6h); means ± SEM of 6 mice. **(h-l)** Short-term 2-DG treatment augments the stemness and antitumor immunity of OT-1 cells. **(h)** 1×10^6^ control or 2-DG-treated OT-1 T cells were adoptively transferred into 6 Gy irradiated C57BL/6 mice 7 days after MC38^OVA^ tumor inoculation. **(i)** Ratio of oxygen consumption rate (OCR) to extracellular acidification rate (ECAR) in control and 2-DG-treated OT-1 T cells *in vitro*; means ± SEM of 3 mice. **(j)** Relative contribution of glycolysis and mitochondrial respiration to cellular ATP production in control and 2-DG-treated OT-1 T cells; means ± SEM of 3 mice. **(l)** Analysis of MC38^OVA^ growth after transfer of 2-DG-treated or control OT-1 T cells; means ± SEM of 4 mice. *, p<0.05; **, p<0.01; ***, p<0.001 by unpaired or paired Student’s t-test in (a-c), (e), (g), (k) and (i). *, p<0.05 by 2-way ANOVA in (l).

## REFERENCES

1. McLane, L.M., Abdel-Hakeem, M.S. & Wherry, E.J. CD8 T Cell Exhaustion During Chronic Viral Infection and Cancer. Annu Rev Immunol 37, 457–495 (2019).

2. Franco, F., Jaccard, A., Romero, P., Yu, Y.R. & Ho, P.C. Metabolic and epigenetic regulation of T-cell exhaustion. Nat Metab 2, 1001–1012 (2020).

3. Williams, M.A. & Bevan, M.J. Effector and memory CTL differentiation. Annu Rev Immunol 25, 171–192 (2007).

4. Zehn, D., Thimme, R., Lugli, E., de Almeida, G.P. & Oxenius, A. ’Stem-like’ precursors are the fount to sustain persistent CD8(+) T cell responses. Nat Immunol 23, 836–847 (2022).

5. Utzschneider, D.T. et al. Early precursor T cells establish and propagate T cell exhaustion in chronic infection. Nat Immunol 21, 1256–1266 (2020).

6. Im, S.J. et al. Defining CD8+ T cells that provide the proliferative burst after PD-1 therapy. Nature 537, 417–421 (2016).

7. Kallies, A., Zehn, D. & Utzschneider, D.T. Precursor exhausted T cells: key to successful immunotherapy? Nat Rev Immunol 20, 128–136 (2020).

8. Utzschneider, D.T. et al. T Cell Factor 1-Expressing Memory-like CD8(+) T Cells Sustain the Immune Response to Chronic Viral Infections. Immunity 45, 415–427 (2016).

9. He, R. et al. Follicular CXCR5-expressing CD8(+) T cells curtail chronic viral infection. Nature 537, 412–428 (2016).

10. Tsui, C. et al. MYB orchestrates T cell exhaustion and response to checkpoint inhibition. Nature (2022).

11. Beltra, J.C. et al. Developmental Relationships of Four Exhausted CD8(+) T Cell Subsets Reveals Underlying Transcriptional and Epigenetic Landscape Control Mechanisms. Immunity 52, 825–841 e828 (2020).

12. Miller, B.C. et al. Subsets of exhausted CD8(+) T cells differentially mediate tumor control and respond to checkpoint blockade. Nat Immunol 20, 326–336 (2019).

13. Siddiqui, I. et al. Intratumoral Tcf1(+)PD-1(+)CD8(+) T Cells with Stem-like Properties Promote Tumor Control in Response to Vaccination and Checkpoint Blockade Immunotherapy. Immunity 50, 195–211 e110 (2019).

14. Wu, T., et al. The TCF1-Bcl6 axis counteracts type I interferon to repress exhaustion and maintain T cell stemness. Sci Immunol 1 (2016).

15. Buck, M.D., Sowell, R.T., Kaech, S.M. & Pearce, E.L. Metabolic Instruction of Immunity. Cell 169, 570–586 (2017).

16. O’Neill, L.A., Kishton, R.J. & Rathmell, J. A guide to immunometabolism for immunologists. Nat Rev Immunol 16, 553–565 (2016).

17. Vaeth, M., Kahlfuss, S. & Feske, S. CRAC Channels and Calcium Signaling in T Cell-Mediated Immunity. Trends Immunol (2020).

18. Vaeth, M. & Feske, S. NFAT control of immune function: New Frontiers for an Abiding Trooper. F1000Res 7, 260 (2018).

19. Wang, Y., Tao, A., Vaeth, M. & Feske, S. Calcium regulation of T cell metabolism. Curr Opin Physiol 17, 207–223 (2020).

20. Vaeth, M. et al. Store-Operated Ca(2+) Entry Controls Clonal Expansion of T Cells through Metabolic Reprogramming. Immunity 47, 664–679 e666 (2017).

21. Klein-Hessling, S. et al. NFATc1 controls the cytotoxicity of CD8(+) T cells. Nat Commun 8, 511 (2017).

22. Shen, H. & Shi, L.Z. Metabolic regulation of TH17 cells. Mol Immunol 109, 81–87 (2019).

23. Scharping, N.E. et al. Mitochondrial stress induced by continuous stimulation under hypoxia rapidly drives T cell exhaustion. Nat Immunol 22, 205–215 (2021).

24. Vardhana, S.A. et al. Impaired mitochondrial oxidative phosphorylation limits the self-renewal of T cells exposed to persistent antigen. Nat Immunol 21, 1022–1033 (2020).

25. Gabriel, S.S. et al. Transforming growth factor-beta-regulated mTOR activity preserves cellular metabolism to maintain long-term T cell responses in chronic infection. Immunity 54, 1698–1714 e1695 (2021).

26. Bengsch, B. et al. Bioenergetic Insufficiencies Due to Metabolic Alterations Regulated by the Inhibitory Receptor PD-1 Are an Early Driver of CD8(+) T Cell Exhaustion. Immunity 45, 358–373 (2016).

27. Siska, P.J., et al. Mitochondrial dysregulation and glycolytic insufficiency functionally impair CD8 T cells infiltrating human renal cell carcinoma. JCI Insight 2 (2017).

28. Yu, Y.R. et al. Disturbed mitochondrial dynamics in CD8(+) TILs reinforce T cell exhaustion. Nat Immunol 21, 1540–1551 (2020).

29. Scharping, N.E. et al. The Tumor Microenvironment Represses T Cell Mitochondrial Biogenesis to Drive Intratumoral T Cell Metabolic Insufficiency and Dysfunction. Immunity 45, 701–703 (2016).

30. Dahling, S. et al. Type 1 conventional dendritic cells maintain and guide the differentiation of precursors of exhausted T cells in distinct cellular niches. Immunity 55, 656–670 e658 (2022).

31. Marchetti, P., Fovez, Q., Germain, N., Khamari, R. & Kluza, J. Mitochondrial spare respiratory capacity: Mechanisms, regulation, and significance in non-transformed and cancer cells. FASEB J 34, 13106–13124 (2020).

32. Pham, A.H., McCaffery, J.M. & Chan, D.C. Mouse lines with photo-activatable mitochondria to study mitochondrial dynamics. Genesis 50, 833–843 (2012).

33. Hochrein, S.M. et al. The glucose transporter GLUT3 controls T helper 17 cell responses through glycolytic-epigenetic reprogramming. Cell Metab 34, 516–532 e511 (2022).

34. Kwong, J.Q. et al. Genetic deletion of the mitochondrial phosphate carrier desensitizes the mitochondrial permeability transition pore and causes cardiomyopathy. Cell Death Differ 21, 1209–1217 (2014).

35. Seifert, E.L., Ligeti, E., Mayr, J.A., Sondheimer, N. & Hajnoczky, G. The mitochondrial phosphate carrier: Role in oxidative metabolism, calcium handling and mitochondrial disease. Biochem Biophys Res Commun 464, 369–375 (2015).

36. Pircher, H., Burki, K., Lang, R., Hengartner, H. & Zinkernagel, R.M. Tolerance induction in double specific T-cell receptor transgenic mice varies with antigen. Nature 342, 559–561 (1989).

37. Zhou, X., Ramachandran, S., Mann, M. & Popkin, D.L. Role of lymphocytic choriomeningitis virus (LCMV) in understanding viral immunology: past, present and future. Viruses 4, 2650–2669 (2012).

38. Chandel, N.S. NADPH-The Forgotten Reducing Equivalent. Cold Spring Harb Perspect Biol 13 (2021).

39. Pedre, B., Barayeu, U., Ezerina, D. & Dick, T.P. The mechanism of action of N-acetylcysteine (NAC): The emerging role of H2S and sulfane sulfur species. Pharmacol Ther 228, 107916 (2021).

40. Khan, O. et al. TOX transcriptionally and epigenetically programs CD8(+) T cell exhaustion. Nature 571, 211–218 (2019).

41. Alfei, F. et al. TOX reinforces the phenotype and longevity of exhausted T cells in chronic viral infection. Nature 571, 265–269 (2019).

42. Chen, J. et al. NR4A transcription factors limit CAR T cell function in solid tumours. Nature 567, 530–534 (2019).

43. Sena, L.A. et al. Mitochondria are required for antigen-specific T cell activation through reactive oxygen species signaling. Immunity 38, 225–236 (2013).

44. Mak, T.W. et al. Glutathione Primes T Cell Metabolism for Inflammation. Immunity 46, 1089–1090 (2017).

45. Man, K. et al. Transcription Factor IRF4 Promotes CD8(+) T Cell Exhaustion and Limits the Development of Memory-like T Cells during Chronic Infection. Immunity 47, 1129–1141 e1125 (2017).

46. Martinez, G.J. et al. The transcription factor NFAT promotes exhaustion of activated CD8(+) T cells. Immunity 42, 265–278 (2015).

47. Majmundar, A.J., Wong, W.J. & Simon, M.C. Hypoxia-inducible factors and the response to hypoxic stress. Mol Cell 40, 294–309 (2010).

48. Martinez-Reyes, I. & Chandel, N.S. Mitochondrial TCA cycle metabolites control physiology and disease. Nat Commun 11, 102 (2020).

49. Corcoran, S.E. & O’Neill, L.A. HIF1alpha and metabolic reprogramming in inflammation. J Clin Invest 126, 3699–3707 (2016).

50. Anso, E. et al. The mitochondrial respiratory chain is essential for haematopoietic stem cell function. Nat Cell Biol 19, 614–625 (2017).

51. Desdin-Mico, G. et al. T cells with dysfunctional mitochondria induce multimorbidity and premature senescence. Science 368, 1371–1376 (2020).

52. Miska, J. et al. HIF-1alpha Is a Metabolic Switch between Glycolytic-Driven Migration and Oxidative Phosphorylation-Driven Immunosuppression of Tregs in Glioblastoma. Cell Rep 27, 226–237 e224 (2019).

53. Fisicaro, P. et al. Targeting mitochondrial dysfunction can restore antiviral activity of exhausted HBV-specific CD8 T cells in chronic hepatitis B. Nat Med 23, 327–336 (2017).

54. Steinert, E.M., Vasan, K. & Chandel, N.S. Mitochondrial Metabolism Regulation of T Cell-Mediated Immunity. Annu Rev Immunol 39, 395–416 (2021).

55. Knudson, A.G., Jr. Mutation and cancer: statistical study of retinoblastoma. Proc Natl Acad Sci U S A 68, 820–823 (1971).

56. Palazon, A. et al. An HIF-1alpha/VEGF-A Axis in Cytotoxic T Cells Regulates Tumor Progression. Cancer Cell 32, 669–683 e665 (2017).

57. Finlay, D.K. et al. PDK1 regulation of mTOR and hypoxia-inducible factor 1 integrate metabolism and migration of CD8+ T cells. J Exp Med 209, 2441–2453 (2012).

58. Koh, H.S. et al. The HIF-1/glial TIM-3 axis controls inflammation-associated brain damage under hypoxia. Nat Commun 6, 6340 (2015).

59. Liikanen, I. et al. Hypoxia-inducible factor activity promotes antitumor effector function and tissue residency by CD8+ T cells. J Clin Invest 131 (2021).

60. Clever, D. et al. Oxygen Sensing by T Cells Establishes an Immunologically Tolerant Metastatic Niche. Cell 166, 1117–1131 e1114 (2016).

61. Doedens, A.L. et al. Hypoxia-inducible factors enhance the effector responses of CD8(+) T cells to persistent antigen. Nat Immunol 14, 1173–1182 (2013).

62. Phan, A.T. et al. Constitutive Glycolytic Metabolism Supports CD8(+) T Cell Effector Memory Differentiation during Viral Infection. Immunity 45, 1024–1037 (2016).

63. Denko, N.C. Hypoxia, HIF1 and glucose metabolism in the solid tumour. Nat Rev Cancer 8, 705–713 (2008).

64. McGettrick, A.F. & O’Neill, L.A.J. The Role of HIF in Immunity and Inflammation. Cell Metab 32, 524–536 (2020).

65. Dengler, V.L., Galbraith, M. & Espinosa, J.M. Transcriptional regulation by hypoxia inducible factors. Crit Rev Biochem Mol Biol 49, 1–15 (2014).

66. Hu, Y. et al. TGF-beta regulates the stem-like state of PD-1+ TCF-1+ virus-specific CD8 T cells during chronic infection. J Exp Med 219 (2022).

67. Ma, C. et al. TGF-beta promotes stem-like T cells via enforcing their lymphoid tissue retention. J Exp Med 219 (2022).

68. Galletti, G. et al. Two subsets of stem-like CD8(+) memory T cell progenitors with distinct fate commitments in humans. Nat Immunol 21, 1552–1562 (2020).

69. Klein Geltink, R.I., et al. Metabolic conditioning of CD8(+) effector T cells for adoptive cell therapy. Nat Metab 2, 703–716 (2020).

70. Mo, F. et al. An engineered IL-2 partial agonist promotes CD8(+) T cell stemness. Nature 597, 544–548 (2021).

71. Sukumar, M. et al. Inhibiting glycolytic metabolism enhances CD8+ T cell memory and antitumor function. J Clin Invest 123, 4479–4488 (2013).

72. Crompton, J.G. et al. Akt inhibition enhances expansion of potent tumor-specific lymphocytes with memory cell characteristics. Cancer Res 75, 296–305 (2015).

73. Shao, M. et al. Inhibition of Calcium Signaling Prevents Exhaustion and Enhances Anti-Leukemia Efficacy of CAR-T Cells via SOCE-Calcineurin-NFAT and Glycolysis Pathways. Adv Sci (Weinh) 9, e2103508 (2022).

74. Kastenmuller, W. et al. Peripheral prepositioning and local CXCL9 chemokine-mediated guidance orchestrate rapid memory CD8+ T cell responses in the lymph node. Immunity 38, 502–513 (2013).

75. Vaeth, M. et al. Dependence on nuclear factor of activated T-cells (NFAT) levels discriminates conventional T cells from Foxp3+ regulatory T cells. Proc Natl Acad Sci U S A 109, 16258–16263 (2012).

