## Supplementary Figures for "HIF-1α-mediated mitochondrial-glycolytic reprogramming controls the transition of precursor to terminally exhausted T cells"

### Extended Data Figure 1

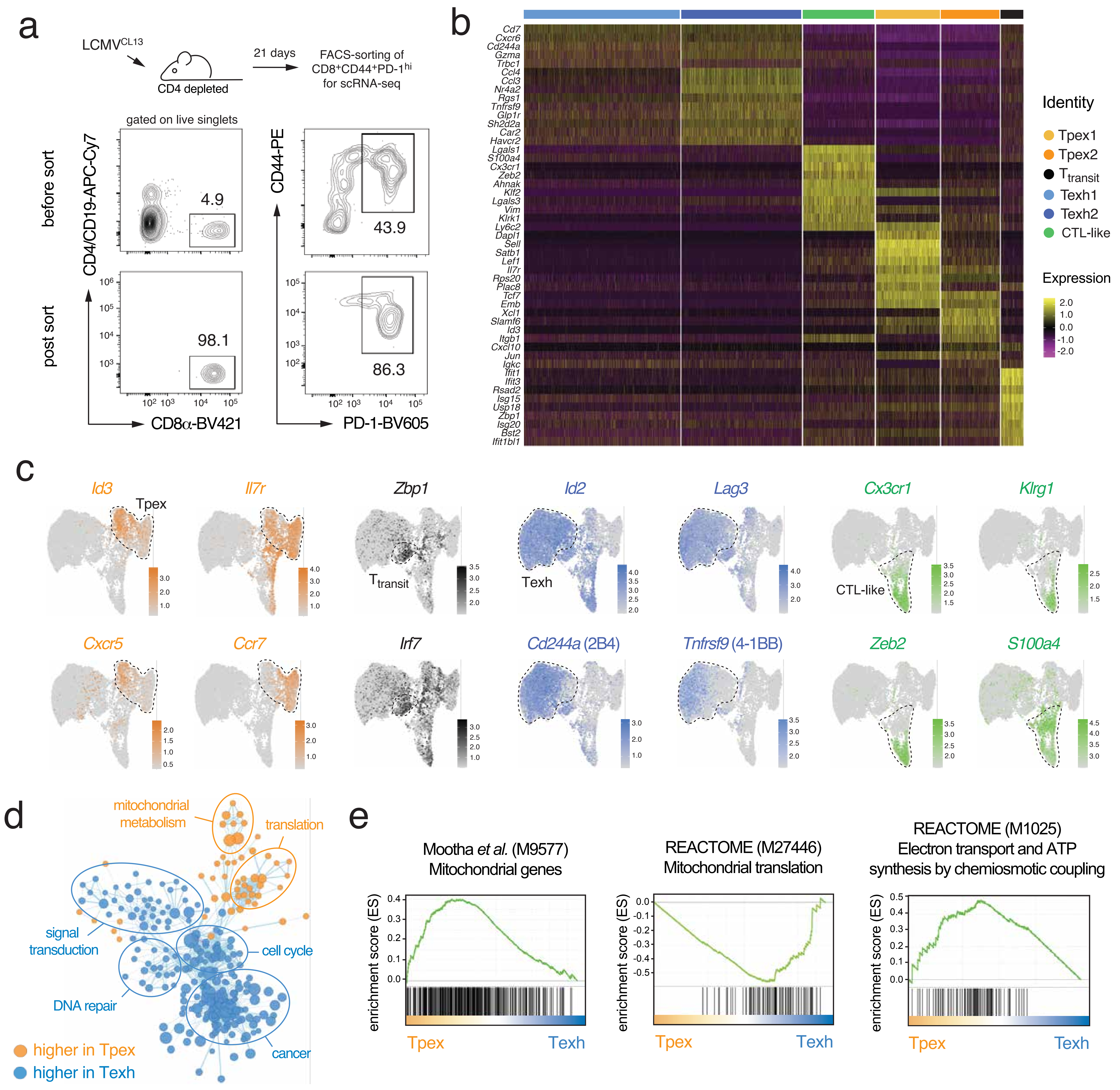

### Extended Data Figure 2

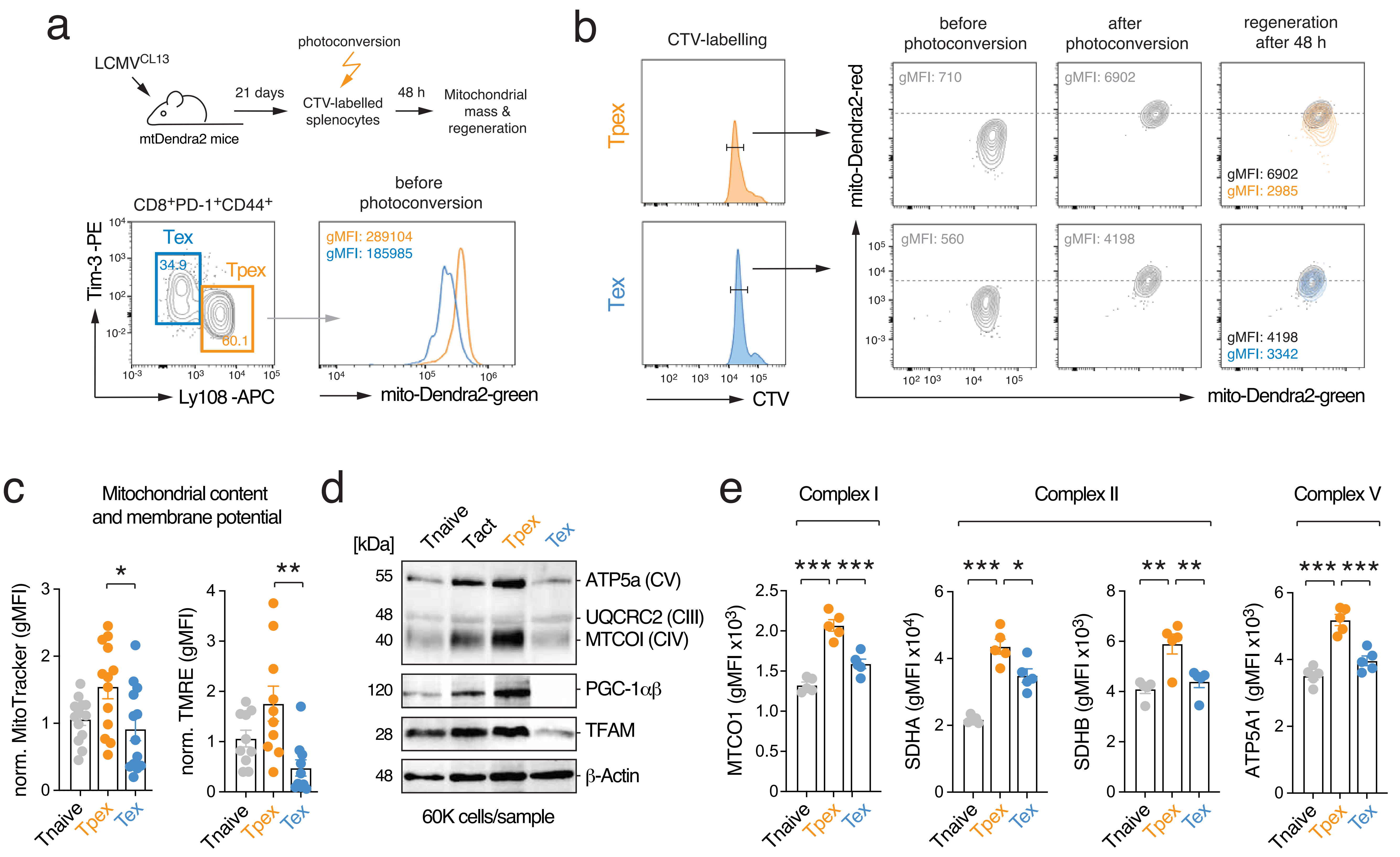

### Extended Data Figure 3

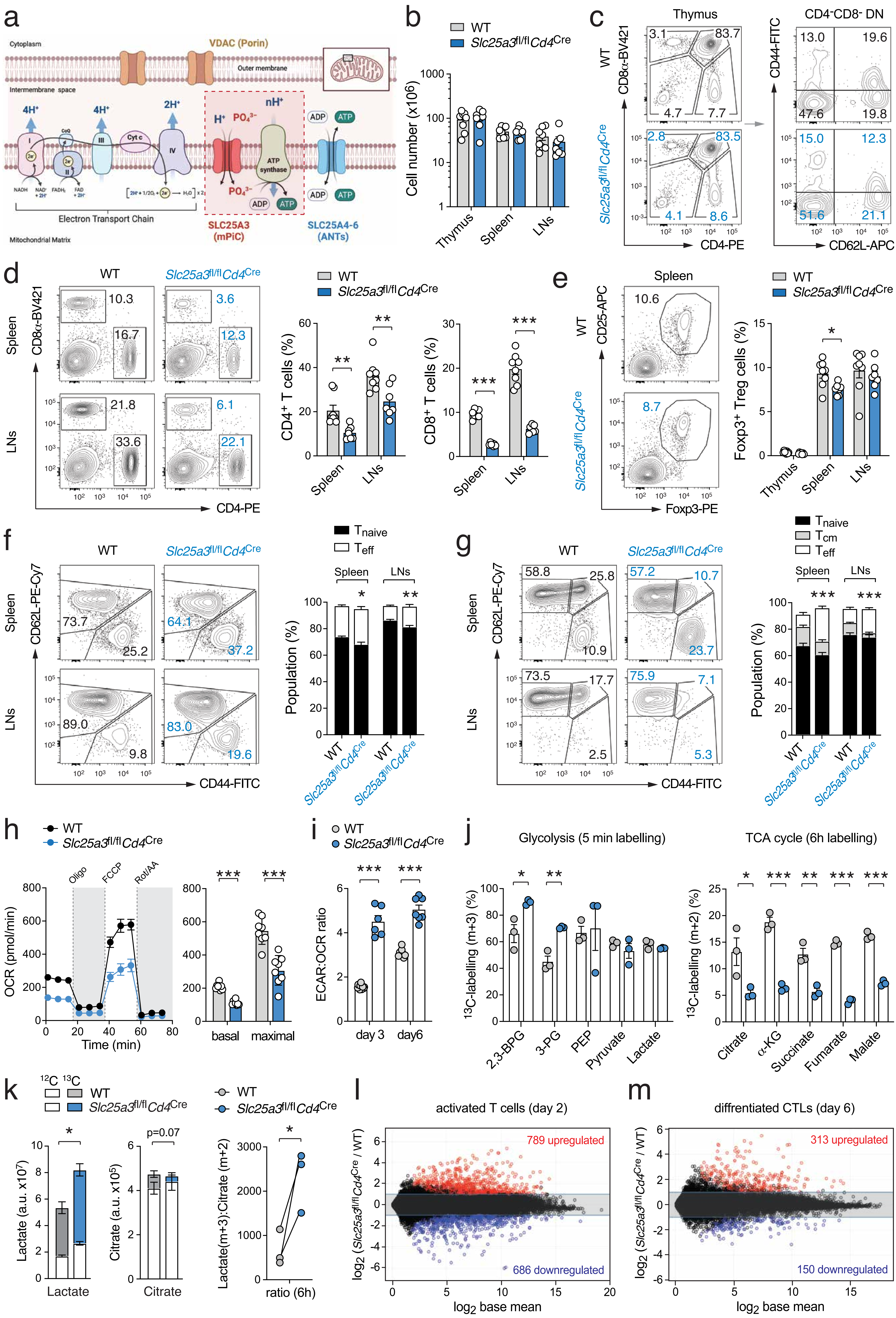

### Extended Data Figure 4

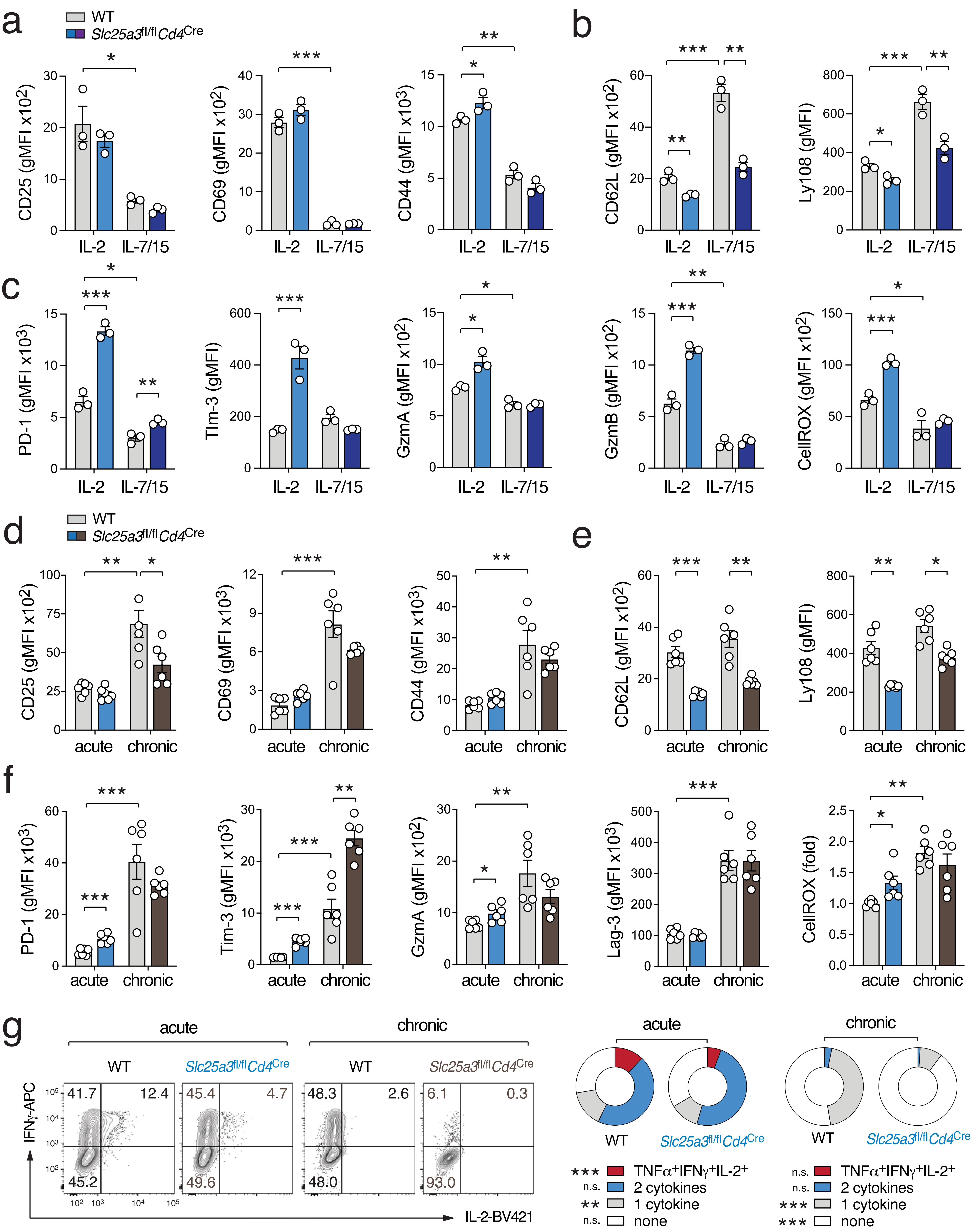

### Extended Data Figure 5

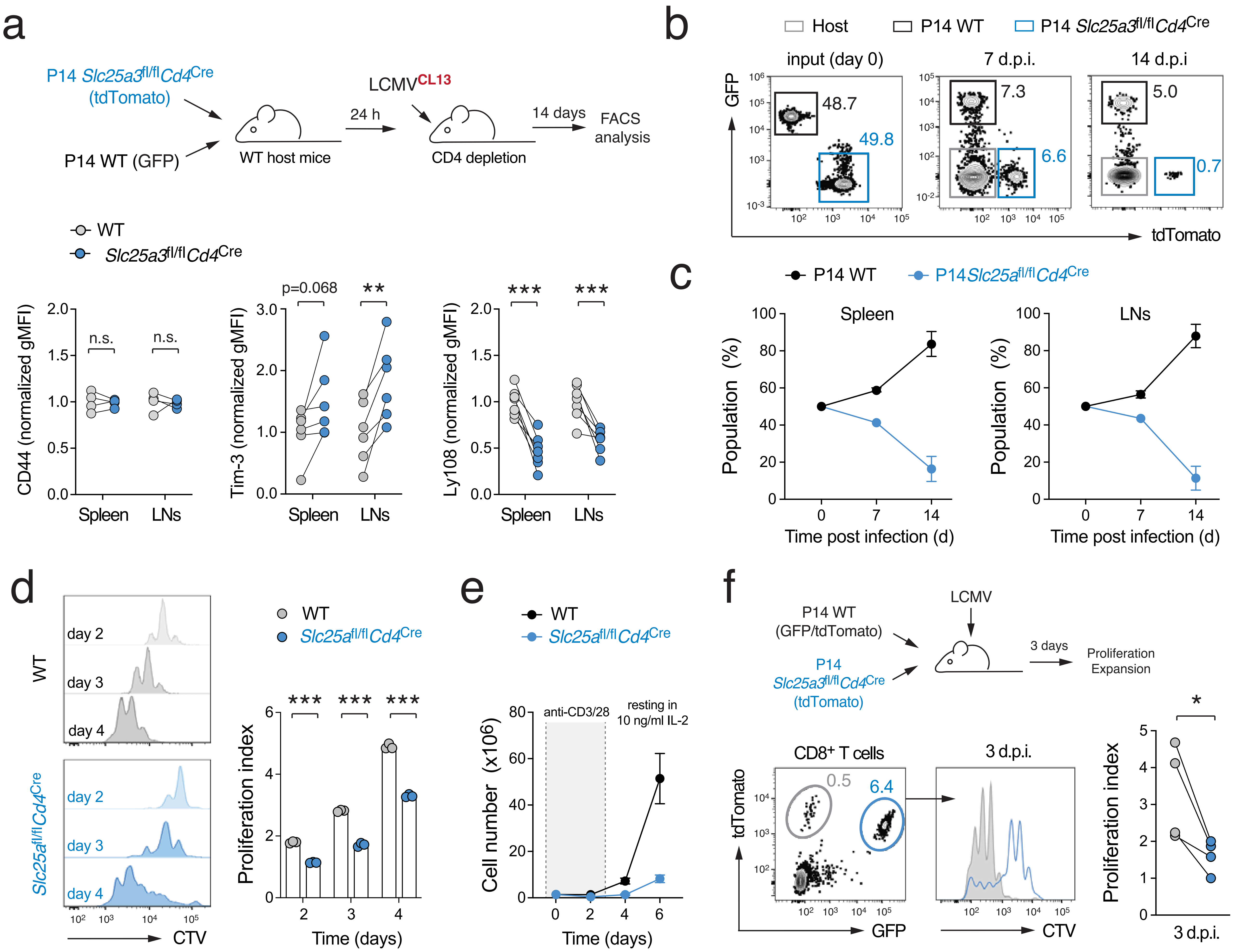

### Extended Data Figure 6

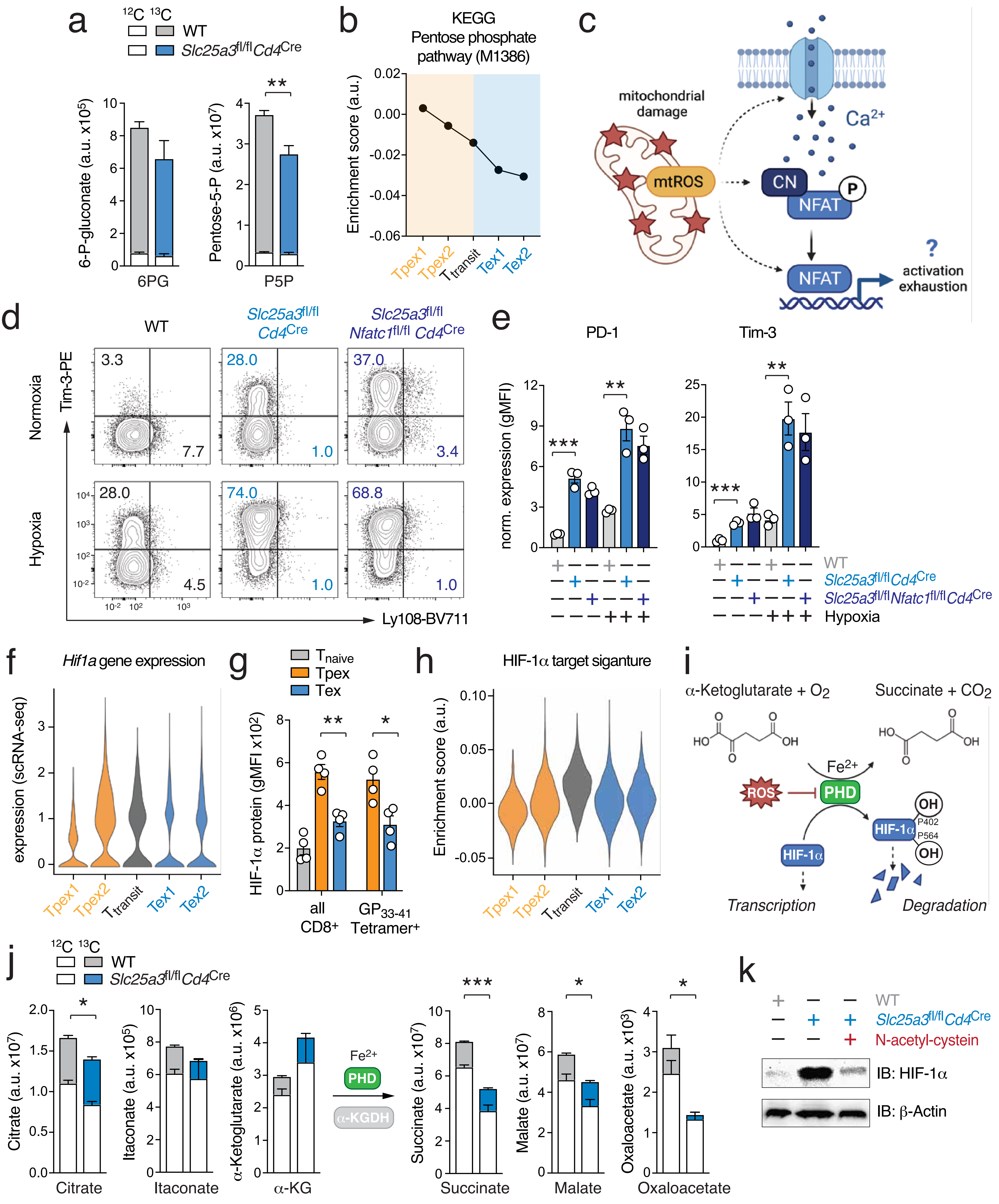

### Extended Data Figure 7

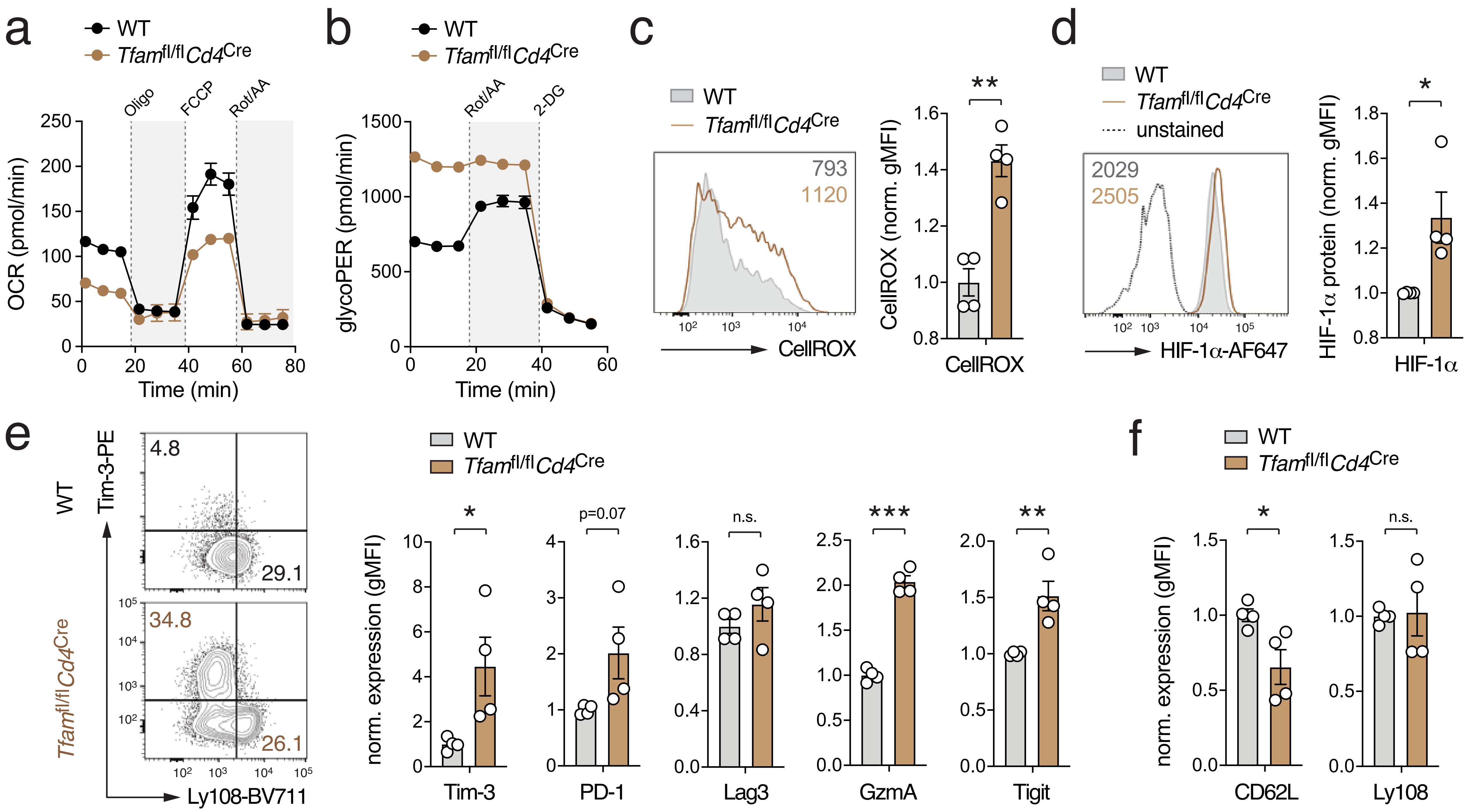

### Extended Data Figure 8

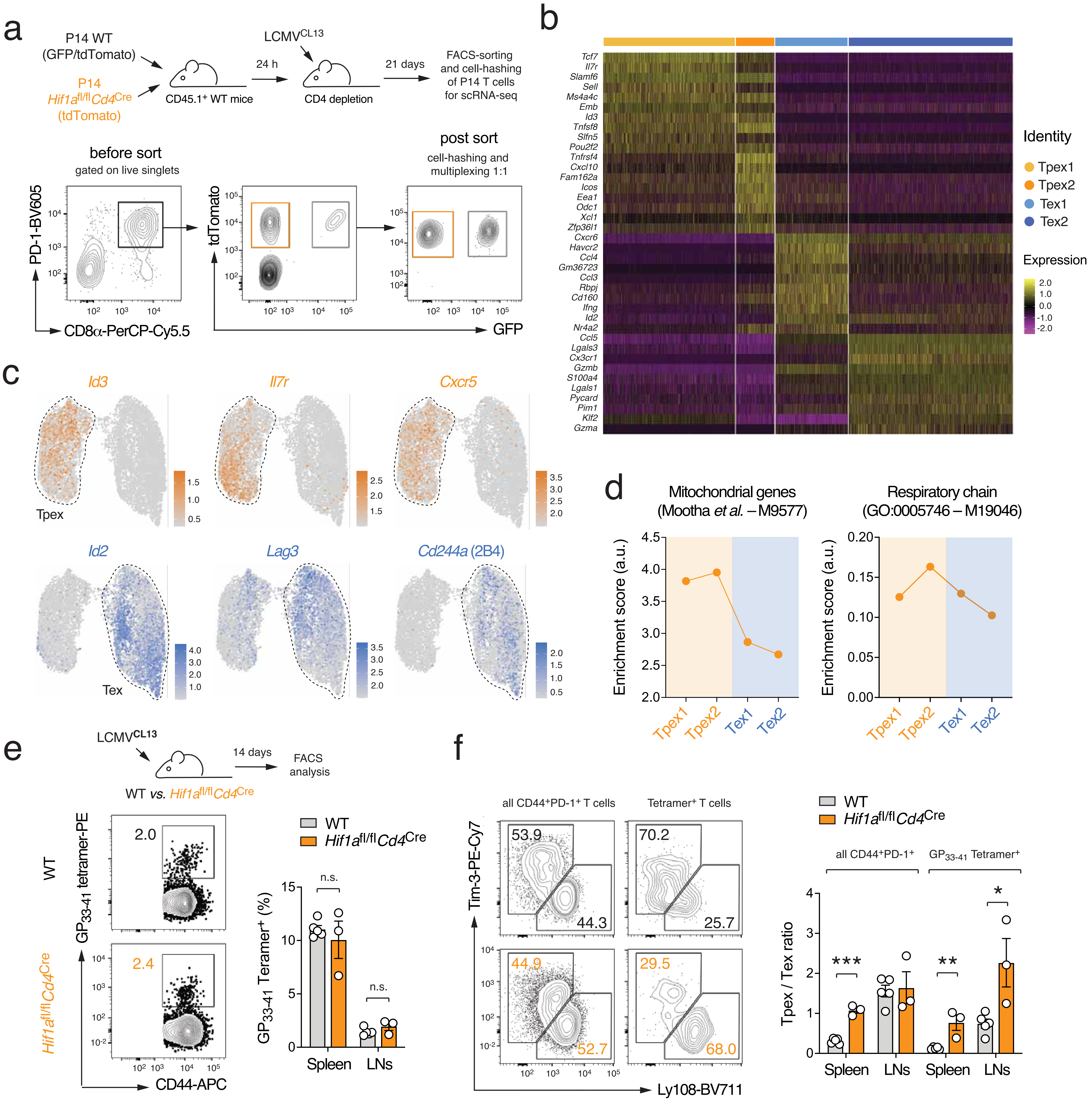

### Extended Data Figure 9

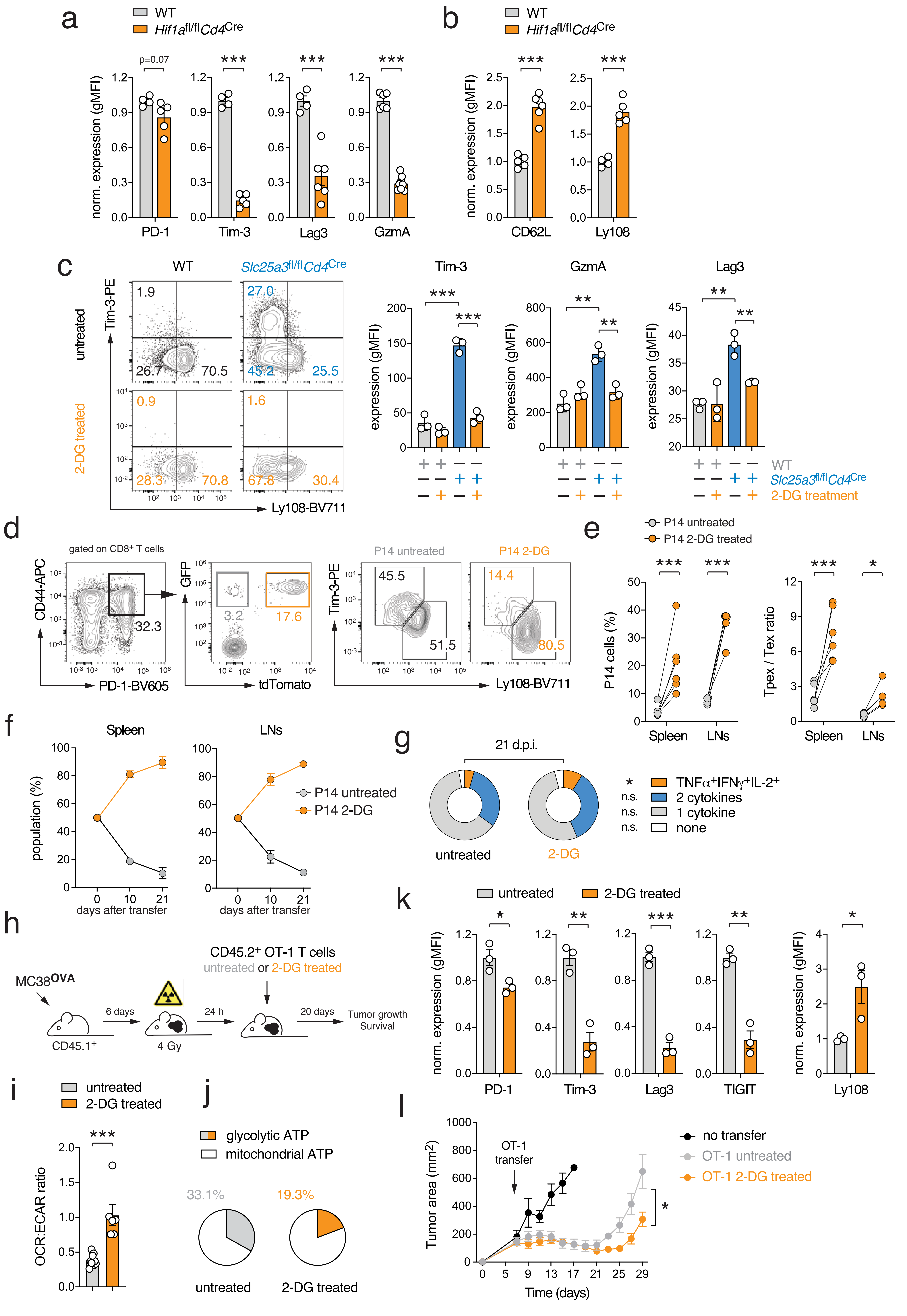
